# Palmitoylation calibrates mitochondrial transport of insulin across the endothelium in insulin resistance

**DOI:** 10.64898/2026.07.14.738288

**Authors:** Wei Zhang, Sangeeta Adak, Qiang Zhang, Gulinu Maimaituxun, Rong Xu, Guifang Dong, Avishek Debnath, Yixuan Wang, Srikanth Singamaneni, Xiaochao Wei, Clay F. Semenkovich

## Abstract

Impaired insulin transport across the endothelium contributes to insulin resistance, a poorly understood condition implicated in diabetes and many other chronic diseases. Insulin has been reported to undergo non-receptor-mediated endocytosis resembling fluid-phase uptake through unclear mechanisms. Here we show in mice with diet-induced insulin resistance that endothelial-specific deletion of the depalmitoylase acyl-protein thioesterase 1 (APT1) improved glucose tolerance and insulin sensitivity without affecting chronic inflammation or capillary structure. Endothelial APT1 deficiency increased interstitial insulin levels in mice confirmed by in-situ microneedle-based sampling. In cultured human microvascular cells, APT1 inhibition enhanced cell transport of high-dose insulin independent of the insulin receptor and canonical endocytic machinery. Unexpectedly, live-cell imaging revealed insulin rapidly localizing to mitochondria prior to endolysosomal trafficking, even at physiological insulin concentrations. APT1 inhibition delayed mitochondrial discharge of insulin to lysosomes. Cyclosporin A, an immunosuppressant known to affect mitochondrial function, preserved mitochondrial insulin content and promoted insulin transport in cultured cells, and enhanced interstitial insulin delivery in mice. Proteomic analysis revealed two palmitoylated proteins, PACS1 and YTHDF2, required for APT1-mediated mitochondrial-endolysosomal trafficking of insulin. A mitochondrial insulin shuttle in endothelial cells may participate in the physiological adaptation to hyperinsulinemia and its regulation by palmitoylation suggests a novel approach to insulin resistance.

## Introduction

Insulin is the dominant mediator of metabolic homeostasis. Defects in insulin signaling causing insulin resistance often precede the onset of type 2 diabetes and related metabolic disorders.^1–3^ Metabolic diseases including myocardial infarction, heart failure, renal failure, Alzheimer’s disease, cancer and others are also linked to insulin resistance, a condition that evades simple mechanistic understanding given its molecular, cellular, and systemic effects.^1–7^

A neglected step in insulin signaling is the transport of insulin from capillaries to the interstitial space. Interstitial insulin is better correlated with glucose disposal than plasma insulin.^8,9^ The insulin gradient between plasma and the interstitium suggests that transendothelial delivery of insulin may be rate-limiting for its action, and that decreased transport across the endothelium may contribute to pathological insulin resistance.^8–14^ Insulin crosses the endothelium through an active process, although the underlying mechanism is controversial.^15,16^ Impaired receptor-mediated signaling in endothelial cells affects glucose metabolism.^17,18^ In contrast, unsaturable, fluid-phase insulin transport has been reported and in skeletal muscle transcapillary transport of insulin in vivo appears to be independent of the insulin receptor.^19,20^ The details of endothelial insulin transport are unknown.^15,16,21^

Mitochondria interact extensively with other cellular compartments, including the endoplasmic reticulum, endosomes, lysosomes, nucleus, and plasma membrane.^22–25^ Mitochondrial dysfunction may occur in insulin resistance.^2,26^ Limited evidence indicates that mitochondria serve as conduits for specific protein cargos in endocytic pathways through interactions with endosomal compartments.^27,28^ Mitochondria are also implicated in an unconventional endocytic process related to protein S-palmitoylation,^29,30^ suggesting communication with the extracellular environment. However, a direct link between mitochondria and insulin transport has not yet been demonstrated.

Protein S-palmitoylation, or S-acylation, is a reversible lipid modification mediated by the actions of acyltransferases, which add palmitate to proteins, and acyl-thioesterases, which remove palmitate.^31,32^ The presence or absence of palmitate allows lipophilic versatility permissive for membrane trafficking. Specific protein palmitoylation events may be involved in diabetes and its complications.^33–38^ Acyl-protein thioesterase 1 (APT1) is a major depalmitoylase associated with both diabetes and mitochondrial function.^39–41^ By studying insulin resistant mice with endothelial APT1 deficiency, we uncovered an unexpected role for mitochondria in insulin delivery across the endothelium.

## Results

### Endothelial APT1 knockout improves glucose tolerance and insulin sensitivity in HFD-fed mice

To study endothelial APT1 deficiency and insulin resistance, we used two previously characterized mouse models: a constitutive Tie2-Cre-mediated APT1 knockout (Tie2-APT1 KO) and an inducible VE-Cadherin-Cre-mediated APT1 knockout (VEC-APT1 KO).^36^ The Tie2-APT1 KO did not alter glucose tolerance in chow diet-fed mice (Figure S1A). High-fat diet (HFD)-fed Tie2-APT1 KO mice had no significant body composition phenotype (Figure 1A, B), but showed improved systemic glucose tolerance and insulin sensitivity compared with littermate controls (Figure 1C, D). HFD-fed VEC-APT1 KO mice also had no body composition phenotype (Figure 1E, F) in the setting of improved glucose tolerance and insulin sensitivity (Figure 1G, H). The phenotype was restricted to male mice, with no detectable changes in females.

**Figure 1.**
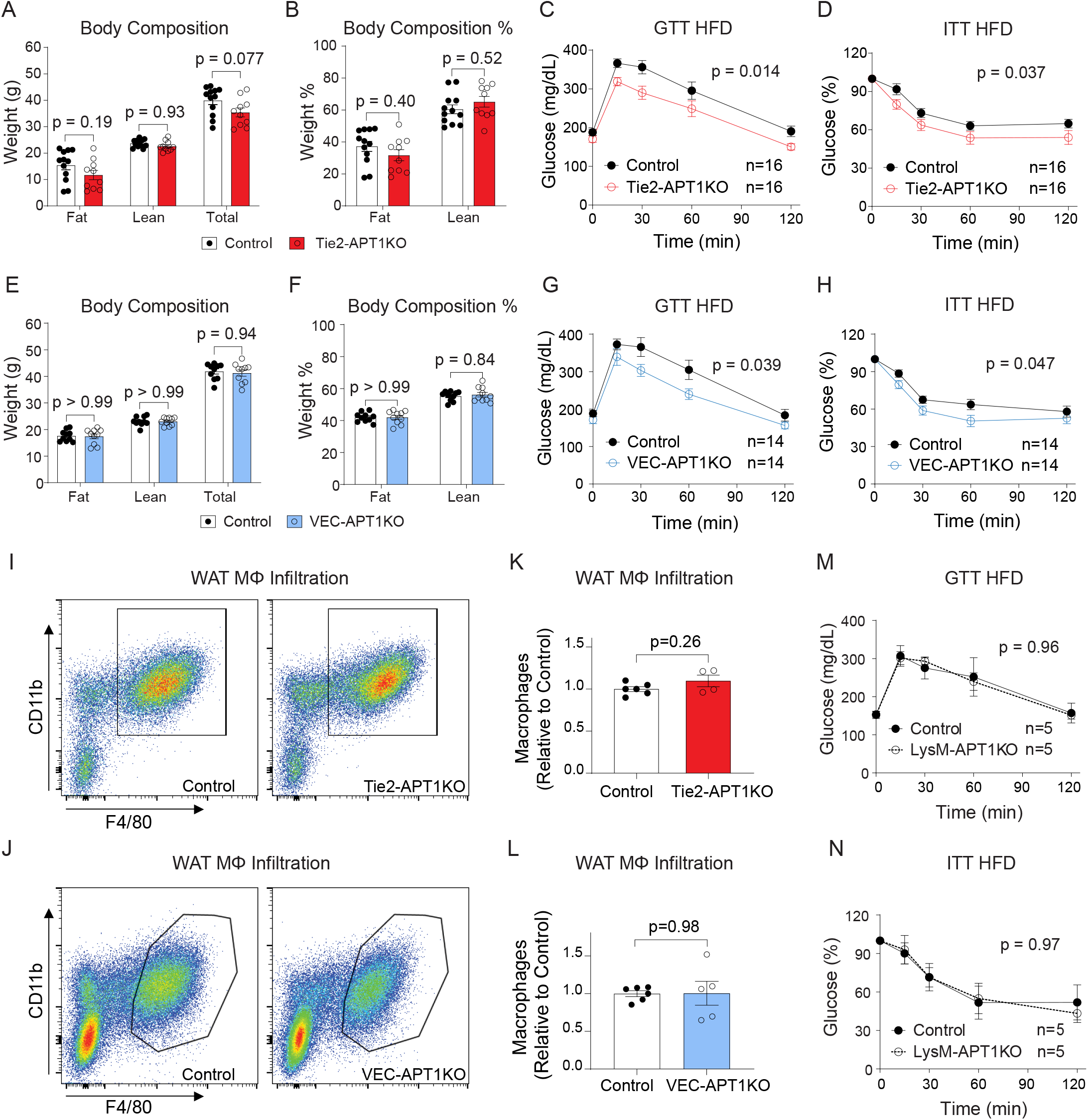
Endothelial-specific APT1 knockout improves glucose tolerance and insulin sensitivity in HFD-fed mice. (**A-D**) Tie2-APT1 KO and littermate control male mice after 3 months of HFD feeding were assessed for body composition (A, B), glucose tolerance tests (GTTs) (C), and insulin tolerance tests (ITTs) (D). (**E-H**) VEC-APT1 KO and littermate control male mice after 3 months of HFD feeding, were assessed for body composition (E, F), GTTs (G), and ITTs (H). (**I-J**) Representative flow cytometry dot plots showing total macrophages (CD11b+, F4/80+) in white adipose tissue of HFD-fed Tie2-APT1 KO mice (I) and VEC-APT1 KO mice (J). (**K-L**) Quantification of adipose tissue macrophage infiltration in HFD-fed Tie2-APT1 KO mice (K) and HFD-fed VEC-APT1 KO mice (L). (**M-N**) GTTs (M) and ITTs (N) for LysM-APT1 KO male mice and littermate control mice after 3 months of HFD feeding. Mice were fasted for 4–6 h prior to GTT or ITT. Data are expressed as mean ± SEM. Statistical analyses for genotype comparisons were performed by two-way ANOVA with Sidak’s post hoc test or by unpaired t-test with Welch’s correction. See also Figure S1.

Since the vasculature mediates inflammation, we assessed immune signatures in adipose tissue, known to be altered in diet-induced insulin resistance.^42,43^ In both models, endothelial APT1 KO did not affect visceral white adipose tissue (WAT) macrophage (CD11b^+^/F4/80^+^) infiltration (Figure 1I, J, K, L, see Figure S1B for gating strategy). Apart from IL-17 in the VEC-APT1 KO model, there was no effect of endothelial APT1 deficiency on inflammatory gene expression (Figure S1C, D). Because Tie2-Cre may induce recombination in hematopoietic cells,^44^ we generated a myeloid cell-specific knockout (LysM-APT1 KO). After HFD feeding, myeloid cell-specific APT1 deletion did not affect glucose tolerance or insulin sensitivity (Figure 1M, N). We also investigated the closely related homolog APT2, which maintains protein palmitoylation homeostasis with APT1.^45^ On HFD, both Tie2-APT2 KO and VEC-APT2 KO mouse models showed no changes in glucose tolerance or insulin sensitivity compared to littermate controls (Figure S1E, F, G, H). Collectively, these data suggest that endothelial APT1 impacts insulin action without major effects on fat mass or inflammation in the setting of HFD feeding.

### Endothelial APT1 deficiency increases interstitial fluid insulin in HFD-fed mice

We utilized a microneedle patch technique^46^ to assess interstitial fluid insulin. With this approach, microneedles coated with capture antibodies specific to human insulin penetrate the epidermis to sample interstitial fluid (ISF). By coupling with an ultrabright plasmonic fluor nanolabel, exogenous insulin content was analyzed on the microneedle patch (Figure 2A). Human insulin (5 IU/kg body weight) was administered by IP (intraperitoneal) injection to HFD-fed control and Tie2-APT1 KO mice then microneedle patches were applied to mouse skin. Fluorescent microneedle signals reflecting exogenous human insulin in ISF were increased in HFD-fed Tie2-APT1 KO mice compared to HFD-fed control mice (Figure 2B, C). To confirm these findings with an alternative approach, skin ISF samples collected by centrifugation from different mouse cohorts also showed increased ISF insulin in HFD-fed Tie2-APT1 KO compared to HFD-fed control mice (Figure 2D). For both approaches, glucose levels were comparable between groups during these acute ISF measurements (Figure S2A, B) despite increased ISF human insulin in KO mice, likely due to the high insulin dose and single time-point analysis. These data suggest that endothelial APT1 deficiency may improve glucose tolerance and insulin sensitivity by increasing transcapillary insulin transport. Consistent with this notion, HFD-fed Tie2-APT1 KO mice exhibited elevated insulin-stimulated AKT phosphorylation in skeletal muscle but not in liver (Figure 2E, F), consistent with structural differences between the continuous endothelium of skeletal muscle and the discontinuous sinusoidal capillaries of the liver.^18^

**Figure 2.**
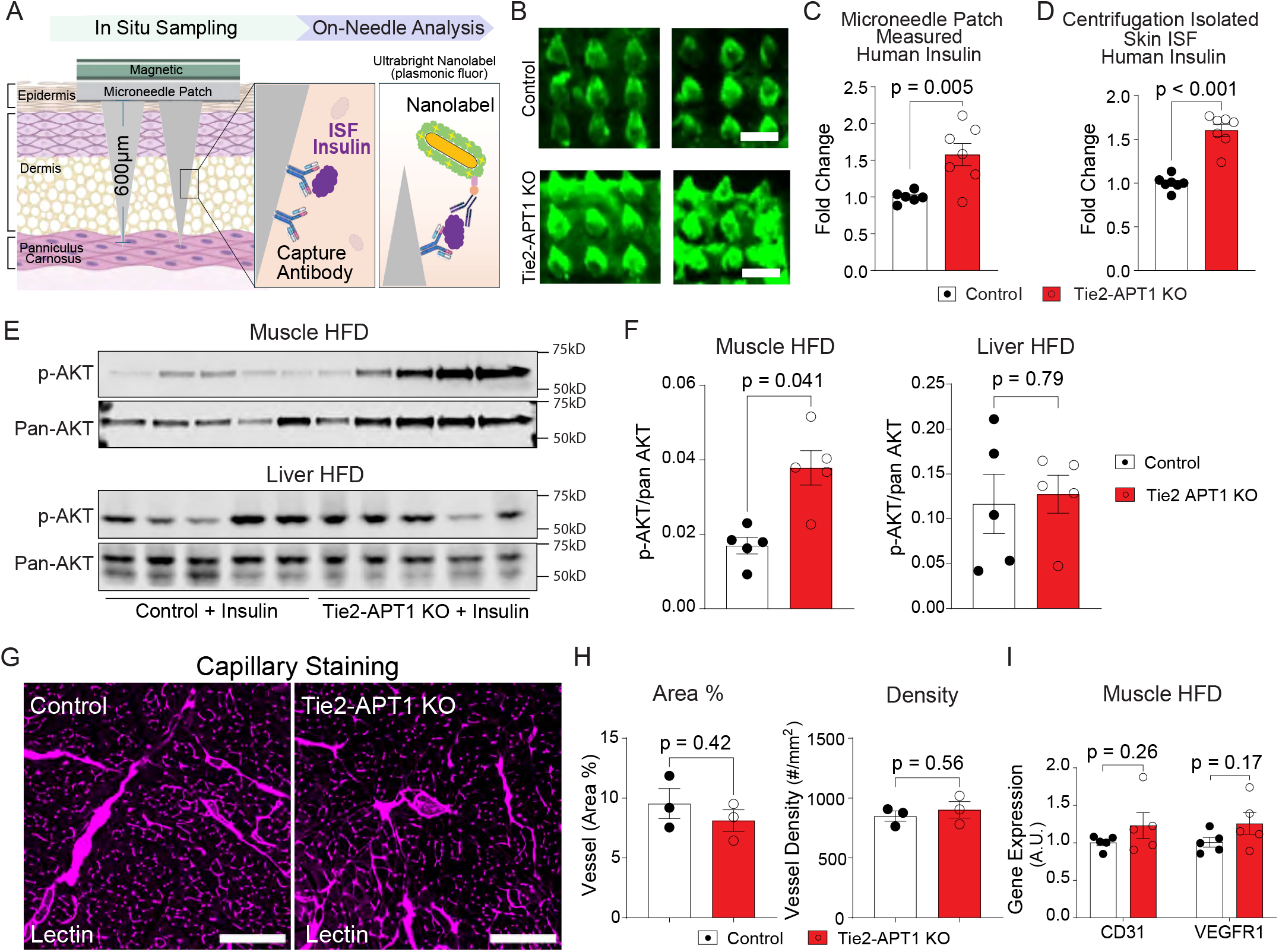
Absence of APT1 in endothelium increases interstitial fluid insulin levels in obese mice. (**A**) Schematic depicting the workflow of microneedle-based immuno-quantitation of interstitial fluid (ISF) insulin level. ISF in situ sampling was achieved with human insulin antibody-coated microneedles applied to the skin, and exogenous human insulin was measured with an ultrabright nanolabel-conjugated detection antibody. (**B**) Representative scanned fluorescence microneedle images from HFD-fed Tie2-APT1 KO mice and littermate control mice (scale bar, 600 μm). Mice were fasted for 5 h before 5 IU/kg human insulin IP-injection. After 10 min, two patches were applied to the abdominal skin for 20 min. (**C**) Quantification of microneedle fluorescence. (**D**) Skin ISF of HFD-fed Tie2-APT1 KO mice and control mice was collected by centrifugation (5 IU/kg human insulin, IP-injection) and human insulin was measured by ELISA. (**E**) Representative Western blots of insulin-induced AKT phosphorylation in skeletal muscle and liver of HFD-fed Tie2-APT1 KO mice and control mice (human insulin 5 IU/kg, 15 min). (**F**) Western blot quantification (p-AKT/pan-AKT ratio) for muscle and liver. (**G-H**) Representative images (G) and quantification (H) of lectin-stained blood vessels in calf muscle. Scale = 200 μm. (**I**) Muscle tissue vessel marker gene expression by q-PCR. Data are expressed as mean ± SEM. Statistical analyses were performed by unpaired t-test or unpaired t-test with Welch’s correction. See also Figure S2.

Increased peripheral tissue vessel density could affect insulin delivery, but immunofluorescent staining of lower limb muscle (Figure 2G) showed comparable vessel area and vessel density between Tie2-APT1 KO and control mice (Figure 2H), indicating that APT1 deficiency did not affect overall muscle vasculature in HFD-fed mice. Moreover, there was no significant increase in gene expression of vascular markers with APT1 deficiency in muscle or adipose tissue of HFD-fed mice (Figure 2I and Figure S2C, D). Transmission electron microscopy (TEM) images of soleus muscle capillaries revealed classic tight junctions of continuous endothelium with widespread intracellular vesicles (Figure S2E). Vesicles were classified as luminal, abluminal, or intracellular and color-coded (Figure S2F). Total vesicle number (Figure S2G) and vesicle density (Figure S2H) were similar in KO and control groups and there were no differences in subcellular vesicle populations between HFD-fed Tie2-APT1 KO and control mice (Figure S2I, J, K).

### APT1 deficiency promotes insulin transport in cultured endothelial cells

Increased interstitial fluid insulin levels in Tie2-APT1 KO mice suggest that endothelium regulates insulin transport. We investigated endothelial insulin transport using a model in which microvascular cells were subjected to short-term insulin exposure to quantify cellular insulin content (uptake), followed by an insulin-free chase to assess extracellular release.^47^ In cultured human adipose microvascular endothelial cells (HAMECs), siRNA-meditated genetic inhibition of APT1 (APT1 KD, Figure 3A) increased insulin uptake and release (Figure 3B,C). Conversely, APT1 overexpression (APT1 OE, Figure 3D) showed a trend toward decreased insulin uptake (Figure 3E, p = 0.092) and significantly decreased insulin release (Figure 3F, p = 0.024). Treatment with the chemical APT1 inhibitor ML348 increased uptake of native (unlabeled) insulin (Figure 3G) as well as fluorescent Alexa Fluor 647-labeled insulin (AF647, Figure 3H and Figure S3A). AF647 insulin and Cy5 insulin, human insulins with different fluorescent labels, and native human insulin showed similar induction of AKT phosphorylation in HAMECs (Figure 3I).

**Figure 3.**
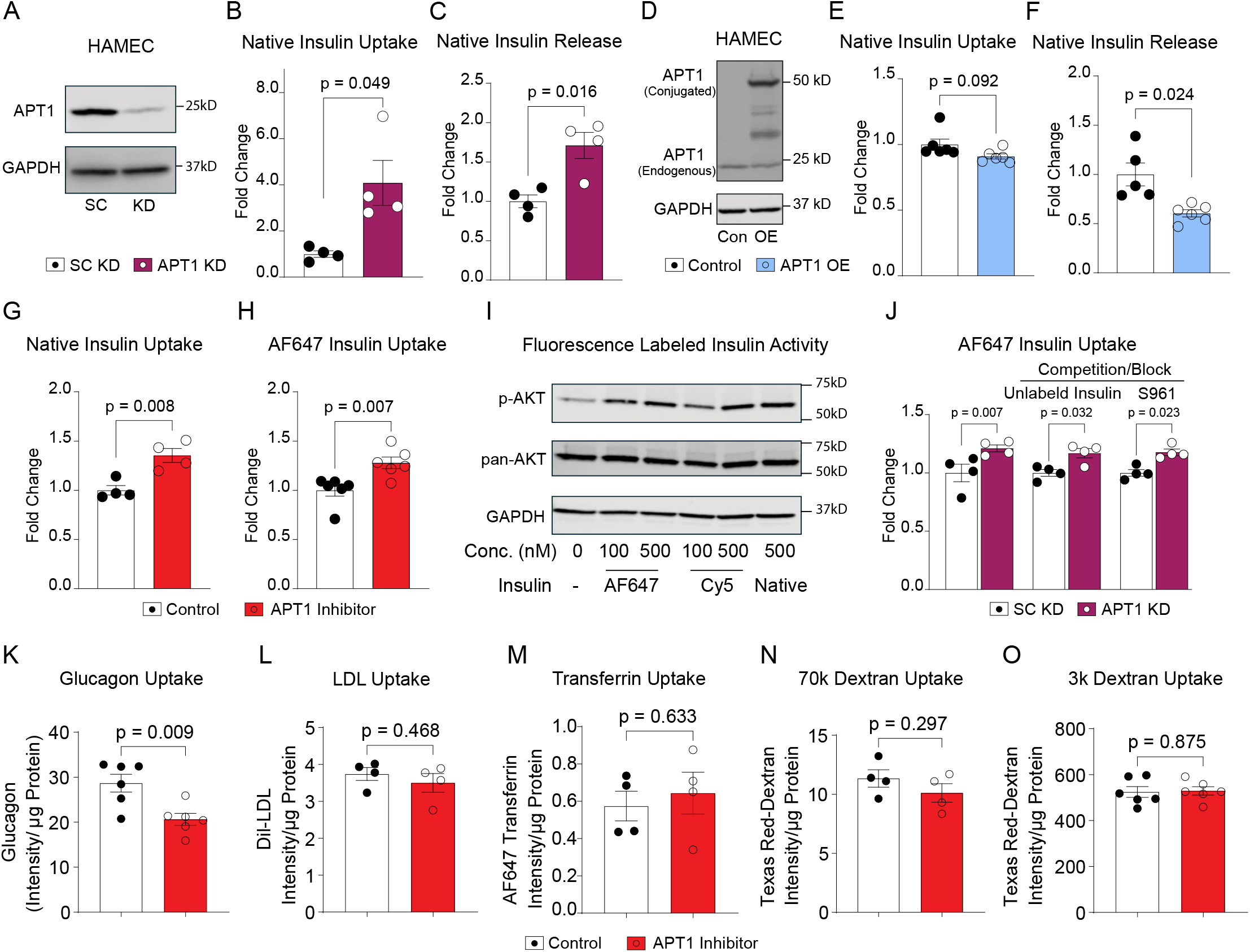
APT1 deficiency promotes insulin uptake in cultured endothelial cells. (**A**) siRNA-mediated APT1 knockdown in HAMECs confirmed by Western blot. (**B-C**) Insulin uptake (B) and insulin release (C) in HAMECs with scrambled or APT1 knockdown. Native insulin treatment: 500 nM for 15 min. Insulin release was collected for 15 min after initial insulin loading. (**D**) Lentivirus-mediated APT1 overexpression in HAMECs confirmed by Western blot. Control: mScarlet only; APT1 OE: mScarlet conjugated APT1 overexpression. (**E**-**F**) Insulin uptake (E) and insulin release (F) in HAMECs with mScarlet control or mScarlet-APT1 overexpression. (**G**) Insulin uptake was assayed in HAMECs loaded with human insulin (500 nM, 15 min) in the absence or presence of the APT1 inhibitor ML348 (10 μM for 3 hrs). (**H**) AF647-labeled insulin uptake in HAMECs treated with or without the APT1 inhibitor ML348. (**I**) Biological activity of native and fluorescent-labeled insulin confirmed by induced AKT phosphorylation in HAMECs. (**J**) AF647 insulin uptake in HAMECs with scrambled control or APT1 knockdown, in the absence or presence of competition (10-fold excess) using unlabeled insulin (5 μM) or insulin receptor antagonist S961 (5 μM) added 5 min prior. (**K**-**O**) Uptake of AF647 glucagon (K, 500 nM), Dil-LDL (L, 5 μg/mL), AF647 transferrin (M, 25 μg/mL), Texas Red-dextran (70kDa) (N, 25 μg/mL), and Texas Red-dextran (3kDa) (O, 25 μg/mL) in HAMECs treated with control or APT1 inhibitor ML348 (10 μM for 3 hrs). Data are expressed as mean ± SEM. Statistical analyses were performed by unpaired t-test with Welch’s correction or two-way ANOVA with Sidak’s post hoc test. See also Figure S3.

Effects of APT1 inhibition on insulin uptake did not appear to require the insulin receptor. Increased uptake of AF647 insulin in APT1 knockdown cells was not affected by inclusion of a 10-fold molar excess of unlabeled insulin or by the presence of the insulin receptor antagonist S961 (Figure 3J). AF647 insulin uptake by human endothelial cells was not inhibited by excess IGF1 (IGF1 compete), excess native insulin (Insulin compete), IGF1 receptor antibody 1H7 (IGF1R Ab), insulin receptor antibody 47-9 (IR Ab), or a combination of IGF1 and insulin receptor antibodies (Figure S3B). Effects of APT1 inhibition on insulin uptake also did not require caveolin or clathrin. Knockdown of clathrin heavy chain (CLTC) or caveolin-1 (Cav1) was confirmed by Western blots (Figure S3C). Neither deficiency of Cav1 or CLTC affected the increased uptake of insulin induced by APT1 inhibition (Figure S3D). Treatment with the clathrin inhibitor Pitstop 2 or methyl-β-cyclodextrin (which depletes membrane cholesterol) did not affect AF647 insulin uptake but uptake was decreased by inhibition of the membrane remodeling protein dynamin (Figure S3E). Unlike insulin (5.7 kD) uptake, glucagon (3.4 kD) uptake was reduced in the presence of APT inhibitor (Figure 3K). Reduced glucagon uptake is unlikely to explain the endothelial-KO phenotype, as glucagon primarily targets the liver, where fenestrated endothelium largely bypasses endothelial transport. The internalization of ∼3,000 kD LDL, ∼76 kD transferrin, 70 kD dextran and 3 kD dextran was unaffected by APT1 inhibition (Figure 3L-O), suggesting that disrupting palmitoylation dynamics does not globally increase uptake of extracellular cargo. These data suggest that APT1 selectively modulates endothelial insulin transport through a non-canonical pathway.

### Endothelial mitochondria are a conduit for insulin transport

Initial imaging of PFA-fixed human endothelial cells (HAMECs) revealed a punctate signal pattern for internalized AF647 insulin. Live-cell imaging was strikingly different, showing tubular structures (Figure S4A left) that were disrupted by PFA fixation to leave punctate signals (Figure S4A right). Live-cell imaging across a broad concentration range revealed prominent tubular structures (Figure 4A). Images shown were independently contrast-adjusted to highlight tubular structures. Increasing laser power and exposure time allowed detection of tubular signals at insulin levels encountered in the human postprandial state (0.5 nM or 500 pM, Figure 4B). No such structures were observed in the control, which displayed only weak autofluorescent speckles. To address the possibility of non-specific dye effects, we treated HAMECs with Cy5-insulin, bearing a different dye. Live-cell imaging of Cy5-insulin showed tubular signals like those of AF647 insulin (Figure S4B), but no tubular structures were observed when cells were treated with AF647 transferrin, a conventional endocytic marker (Figure S4C). Unless otherwise noted, subsequent imaging experiments were performed at 50–100 nM to ensure sufficient signal-to-noise ratio. AF647 insulin did not colocalize with 70 kD dextran or 3 kD dextran, pinocytosis markers (Figure S4D). Imaging studies demonstrated that insulin uptake was temperature-sensitive and dynamin-dependent (Figure S4E, F). Figure 4C shows that tubular structures of AF647 insulin in HAMECs colocalized with mitochondrial markers, including MitoTracker (matrix) and GFP-conjugated markers for the outer membrane (MAO, monoamine oxidase A) and the inner membrane (COX8, cytochrome c oxidase subunit VIII), but not with an ER marker (Figure 4D). In contrast, internalized glucagon did not localize in mitochondria (Figure 4E). Higher resolution live-cell SIM (Structured Illumination Microscopy) confirmed insulin inside mitochondria (Figure 4F). These images suggest that a mitochondrial-associated tubular network may be a previously unrecognized trafficking compartment for insulin in endothelial cells.

**Figure 4.**
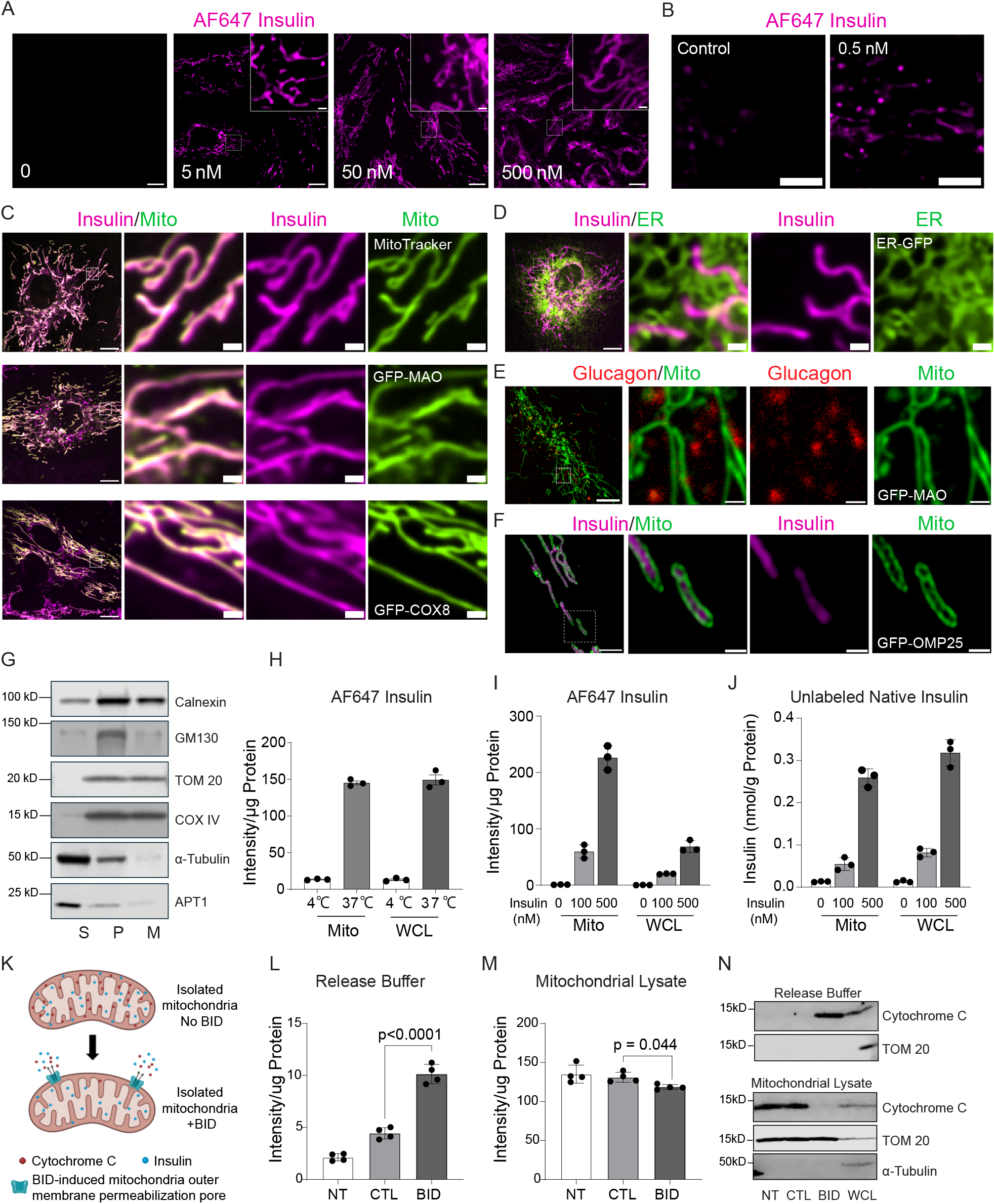
Cellular uptake of insulin leads to mitochondrial localization. (**A**) Representative live-cell confocal images of HAMECs treated with 5 to 500 nM AF647 insulin. Images were independently contrast-adjusted to highlight detailed structures at various insulin dosages, scale = 10 μm (1 μm in inserts). (**B**) Live-cell confocal images of HAMECs treated with 0.5 nM AF647 insulin acquired with increased laser power and exposure time. Control and 0.5nM images with same contrast setting. (B-F) are all from HAMECs with live-cell imaging. (**C**) Localization of AF647 insulin within mitochondria. Cells stained with MitoTracker green FM dye or expressing markers for the mitochondria outer membrane (GFP-MAO), mitochondria inner membrane (GFP-COX8) and loaded with 50 nM AF647 insulin. (**D**) Confocal images were acquired for ER (CellLight ER-GFP) and 50 nM AF647 insulin. (**E**) Confocal images were acquired for glucagon (500 nM AF647 glucagon) with mitochondria (MitoTracker). Scale = 10 μm (1 μm in zoomed-in images). (**F**) Super resolution SIM images were acquired for insulin (500 nM AF647 insulin) with mitochondria outer membrane marker (EGFP-OMP25). Scale = 2 μm (0.5 μm in zoomed-in images). (**G**) Fractions from centrifugation including isolated mitochondria (M), supernatant (S), and pellet (P) were subjected to Western blot analysis. (**H**) AF647 insulin in mitochondria isolated using a biochemical centrifugation approach after insulin loading (500 nM, 15 min) at 4°C or 37°C. Mito: mitochondria, WCL: whole cell lysates. (**I-J**) AF647 insulin (I) and native insulin (J) content in mitochondria isolated from HAMECs treated with vehicle, 100 nM, or 500 nM insulin for 15 min. Mito: mitochondria, WCL: whole cell lysates. (**K**) Schematic of human BID peptide (caspase-8-cleaved)-induced mitochondria outer membrane permeabilization pore formation and cytochrome C release. To confirm the presence of insulin in mitochondria, HAMECs were treated with 500 nM AF647 insulin, then isolated mitochondria were incubated with 100 nM BID peptide for 30min. (**L-M**) BID-induced insulin release (L) and remaining insulin (M) in mitochondria isolated from HAMECs. NT: no treatment, CTL: vehicle control. (**N**) BID activity was confirmed by cytochrome C release from mitochondria into the release buffer by Western blot analysis. Data are expressed as mean ± SEM. Statistical analyses were performed by one-way ANOVA with Dunnett T3 post hoc test. See also Figure S4.

To confirm this finding biochemically, we isolated mitochondria from HAMECs by differential centrifugation with Western blot analysis of fractions shown in Figure 4G. Internalized AF647 insulin was recovered in mitochondrial fractions, which requires physiological temperature (Figure 4H). Both labeled (Figure 4I) and native (Figure 4J) insulin were enriched in mitochondrial fractions in a dose-dependent manner in HAMECs. Caspase-8-cleaved BID peptides permeabilize the mitochondrial outer membrane while preserving mitochondrial matrix content (see schematic in Figure 4K).^48^ As shown in Figure 4L and 4M, BID treatment increased insulin released from isolated mitochondria, accompanied by a modest decrease of insulin remaining in the mitochondria fraction. These findings support a strong association between insulin and mitochondria, with most insulin contained in the matrix. Successful BID-induced outer membrane permeabilization was confirmed by the release of cytochrome C into the release buffer (Figure 4N). TEM images of skeletal muscle capillary endothelium showed mitochondria adjacent to membrane-invaginated vesicles (Figure S4G) close to the plasma membrane (Figure S4H), and even protruding into the capillary lumen (Figure S4I). Additionally, live-cell TIRF imaging of HAMECs revealed that both insulin and mitochondria moved within the thin evanescent zone (Figure S4J, K) close to the cell surface. These data together suggest that insulin uptake may involve a temporal and spatial interaction between mitochondria and the plasma membrane.^49^

### Reduced mitochondrial insulin egress to late endosomal vesicles is associated with enhanced uptake

Besides mitochondrial localization, internalized insulin also appeared as vesicular signals. Characterization with endosome/lysosome markers showed that vesicular insulin partially localized to Rab7a^+^ late endosomes and Lamp1^+^ lysosomes, but not to Rab5a^+^ early endosomes or Rab11b^+^ recycling endosomes (Figure 5A). We next examined the dynamics of intracellular insulin trafficking with a cold-start experiment. As shown in Figure S5A (protocol on top, images over time below), HAMECs were loaded with AF647 insulin at 4° C, which suppresses uptake, followed by time-lapse imaging as the temperature increased. The tubular network was predominant at time 0, suggesting that mitochondria constitute the primary site for initial insulin uptake. With increasing temperature, insulin gradually appeared in apparent vesicles. We noticed an apparent loss of mitochondrial signal that was different from prior experiments, likely due to significantly weak signal resulting from suppressed uptake under the “cold-start” conditions. This transfer from mitochondria to vesicles was studied in HAMECs expressing a mitochondrial marker, where imaging showed insulin rapidly exiting mitochondria into peri-mitochondrial vesicles (arrows in Figure 5B, Video S1). This finding was consistent with time-lapse imaging of HAMECs expressing GFP-Rab7a (Video S2), which shows insulin exiting from tubules and entering Rab7a-marked vesicles (arrows in Figure 5C).

**Figure 5.**
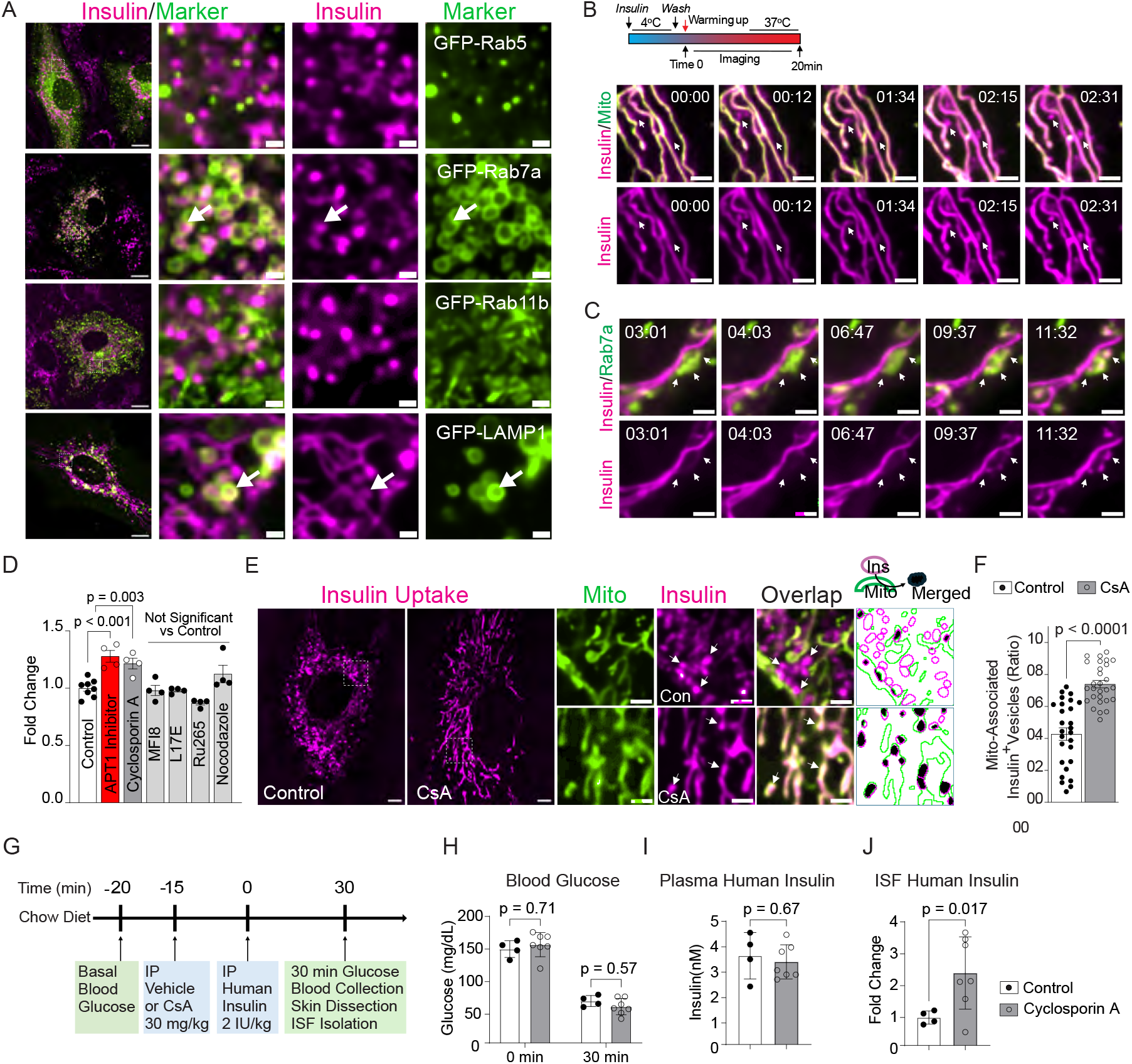
A mitochondrial-to-late-endosome trafficking axis attenuates insulin uptake. **(A)** Co-imaging of AF647 insulin and endo-lysosomal markers. Scale = 10 μm (1 μm for zoomed-in images). **(B)** Representative time-lapse images of AF647 insulin and a mitochondrial marker (GFP-COX8) in HAMECs with timestamps (mm:ss) indicating time after the imaging chamber was warmed up. A schematic of the insulin loading, washing, and imaging steps is shown. Scale = 2 μm. (**C**) Time-lapse images of AF647 insulin with the late endosome marker Rab7a in HAMECs, with timestamps (mm:ss) and schematics as described above. Scale = 2 μm. (**D**) AF647 insulin uptake in HAMECs treated with control buffer, APT1 inhibitor (10 μM), mPTP inhibitor Cyclosprin A (CsA, 5 μM), mitofusin 2 inhibitor MFI8 (10 μM), late lysosome disruption reagent L17E (40 μM), mitochondrial calcium uniporter inhibitor Ru265 (50 μM), or microtubule assembly inhibitor Nocodazole (3 μM). (**E**) Representative live-cell images of AF647 insulin and a mitochondrial marker (GFP-MAO) in HAMECs treated with control or CsA (5 μM). The last column shows quantified areas of mitochondria (green) and vesicular insulin (purple), with regions of colocalization highlighted in black. Scale = 5 μm (2 μm for zoomed-in images). (**F**) Quantification of mitochondria-associated insulin^+^ vesicles in control and CsA-treated cells. (**G**) Schematic diagram of CsA experiment in mice. (**H**) Blood glucose before and after human insulin injection. (**I**) Plasma human insulin 30 min after insulin injection. (**J**) Skin ISF insulin of wildtype mice treated with vehicle control or CsA. Data are expressed as mean ± SEM. Statistical analyses were performed by unpaired t test with Welch’s correction, one-way ANOVA or two-way ANOVA with Sidak’s post hoc test. See also Video S1-2 and Figure S5.

To assess mitochondria-to-late endosome insulin trafficking, we studied effects of mitochondria and endosome inhibitors on insulin uptake. As shown in Figure 5D, similar to APT1 inhibition, cyclosporin A (CsA) significantly increased insulin uptake in HAMECs, but targeting mitofusin1/2 (MFI8), late endosomes (L17E), the calcium uniporter (Ru265), or microtubule assembly (Nocodazole) had no effect. CsA limits opening of the mitochondrial permeability transition pore (mPTP) to impact mitochondrial function,^50^ a mechanism potentially relevant to insulin transport. Under live-cell imaging, CsA-treated cells showed prolonged association of insulin vesicles with mitochondria (Figure 5E, F), implicating mitochondrial function in insulin transport across endothelial cells, a process likely opposed by the endolysosomal pathway. We also tested the effect of CsA on ISF insulin in vivo (Figure 5G). In chow diet-fed mice given human insulin, glucose responses and plasma insulin levels were unchanged after CsA treatment (Figure 5H, I), but CsA, similar to APT1 inhibition in HFD-fed animals, increased insulin levels in skin ISF (Figure 5J).

### APT1 inhibition reinforces mitochondrial retention of insulin through vesicular sequestration

We next examined whether the effect of APT1 on insulin uptake involves mitochondrial-to-late-endosome transition. APT1 inhibition increased mitochondrial insulin levels, demonstrated by both biochemical fractionation (Figure 6A) and TOM22 bead-mediated pull-down (Figure 6B, with Western blot analysis of fractions shown in Figure S6A). APT1 inhibition did not appear to affect the overall morphology of mitochondria (Figure S6B) or Rab7a^+^ endosomes (Figure S6C). Rather, APT1 deficiency (KD confirmed in Figure S6D) significantly decreased the motility of insulin-containing vesicles at physiological temperature (Video S3, Figure 6C-E), implying an extended association of insulin with mitochondria. This implication was validated by imaging of HAMECs with GFP-labeled mitochondria at 37° C (Figure 6F) demonstrating that APT1 inhibition significantly promoted the retention of insulin-containing vesicles at mitochondria (Figure 6G).

**Figure 6.**
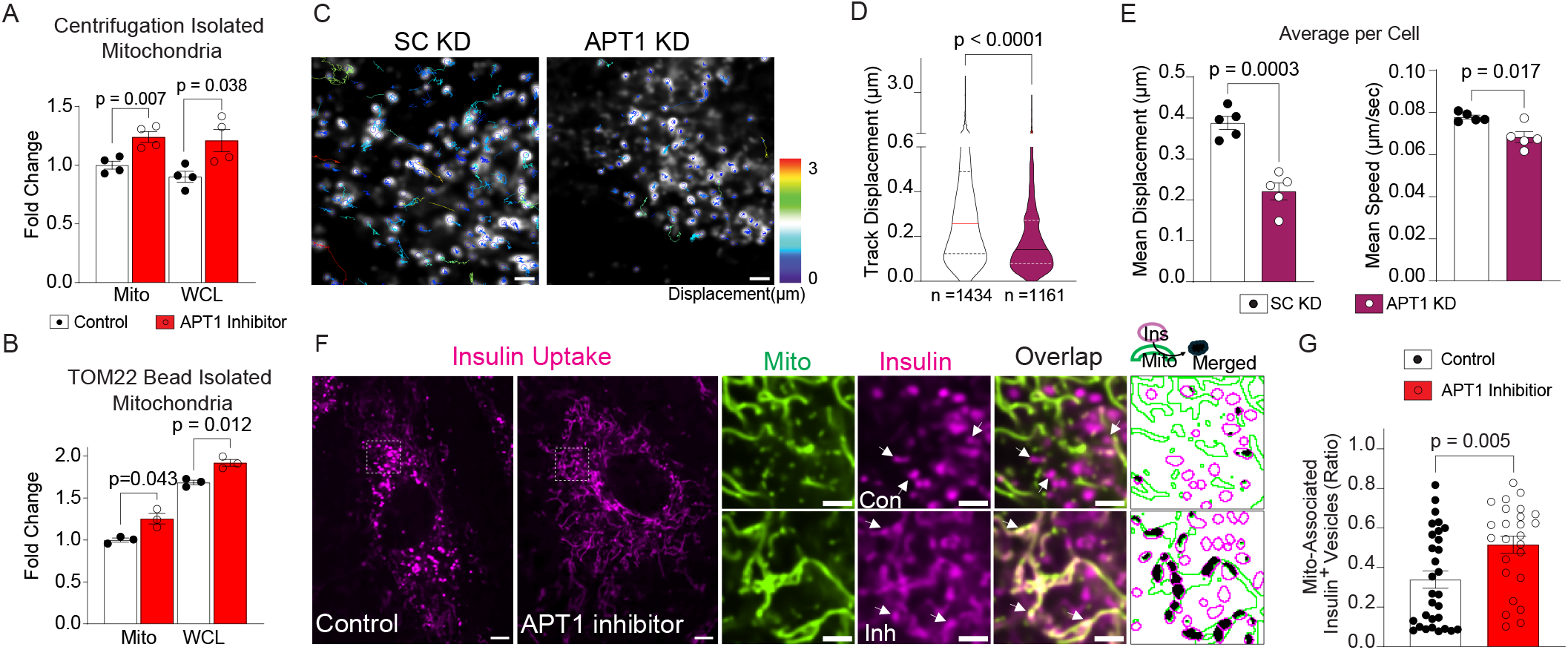
APT1 inhibition reinforces mitochondrial insulin retention via vesicular sequestration. (**A-B**) APT1 inhibition increased insulin content measured in isolated mitochondria by centrifugation method (A) and TOM22 beads (B). Mito: mitochondria. WCL: whole cell lysates. (**C-E**) Tracking analysis of insulin-positive vesicles in HAMECs treated with scrambled control or APT1 knockdown. Representative live-cell images of AF647 insulin marked vesicles with overlaid color-coded tracking trajectories indicating displacement (C). Scale = 2 μm. (D) Displacement of individual vesicle tracks. (E) Average track displacement and mean speed for each cell. (**F**) Representative live-cell images of AF647 insulin and a mitochondrial marker (GFP-MAO) in HAMECs with or without APT1 inhibition, acquired at 37°C. The last column shows quantified areas of mitochondria (green) and vesicular insulin (purple), with regions of colocalization highlighted in black. Scale = 5 μm (2 μm for zoomed-in images). (**G**) Quantification of mitochondria-associated insulin^+^ vesicles in control and APT1-inhibited cells. Statistical analyses were performed by unpaired t test with Welch’s correction. See also Figure S6.

### PACS1 and YTHDF2 as mediators of APT1-regulated insulin transport

We used palmitoylation proteomics to search for potential substrates mediating the effect of APT1 on insulin transport. Three pairs of APT1 KD and scrambled control (SC) endothelial cells, treated with no insulin (basal), 10 nM insulin, or 500 nM insulin, were used to purify palmitoylated proteins by acyl resin-assisted capture (acyl-RAC) followed by mass spectrometry (Figure 7A). APT1 knockdown in each pair was confirmed by Western blot (Figure S7A), acyl-RAC enrichment was confirmed by gel staining (Figure S7B), and principal component analysis (PCA) confirmed robust clustering of the experimental groups (Figure S7C). Heatmap and volcano plots of putative palmitoylated proteins are shown in Figure S7D, E. Using thresholds of unadjusted p < 0.05, ≥ 2 unique peptides, and fold change ≥ 1.2, we identified 979 potential APT1 substrate proteins in HAMECs across all three treatment pairs. There are 24 candidates common to all three groups (Figure 7B). Based on effect size and functional relevance, we selected PACS1 (phosphofurin acidic cluster sorting protein 1), YTHDF2 (YTH N6-methyladenosine RNA binding protein 2), and STX12 (Syntaxin12) for further study (quantitative analyses are shown in Figure S7F). PACS1 and STX12 are involved in vesicular trafficking, and YTHDF2 is associated with mitochondria function. APT1-dependent palmitoylation was validated for all three candidates by acyl-RAC assay (Figure 7C). Knockdown (validation shown in Figure S7G) of PACS1 or YTHDF2, but not Syntaxin 12, abolished the effect of APT1 inhibition on AF647 insulin uptake (Figure 7D). Palmitoylation of PACS1 and YTHDF2 proteins was further confirmed using a metabolic labeling assay combining palmitate-alkyne incorporation with Cy5.5-azide click chemistry detection (Figure 7E). Knockdown of PACS1 or YTHDF2 in HAMECs abolished the effect of APT1 inhibition on AF647 insulin release (Figure 7F, G). Analysis of mitochondrial trafficking of AF647 insulin in knockdown cells showed that in contrast to scrambled control cells, APT1 inhibition failed to increase the association of insulin vesicles with mitochondria in PACS1-or YTHDF2-depleted cells (images in Figure 7H, knockdown confirmed in Figure 7I, quantification in Figure 7J). These results support a model in which PACS1 and YTHDF2 depalmitoylation is required for mitochondrial insulin discharge. APT1 inhibition leads to accumulation of palmitoylated PACS1 and YTHDF2 that cooperatively drive mitochondrial insulin retention and promote insulin transport in endothelial cells.

**Figure 7.**
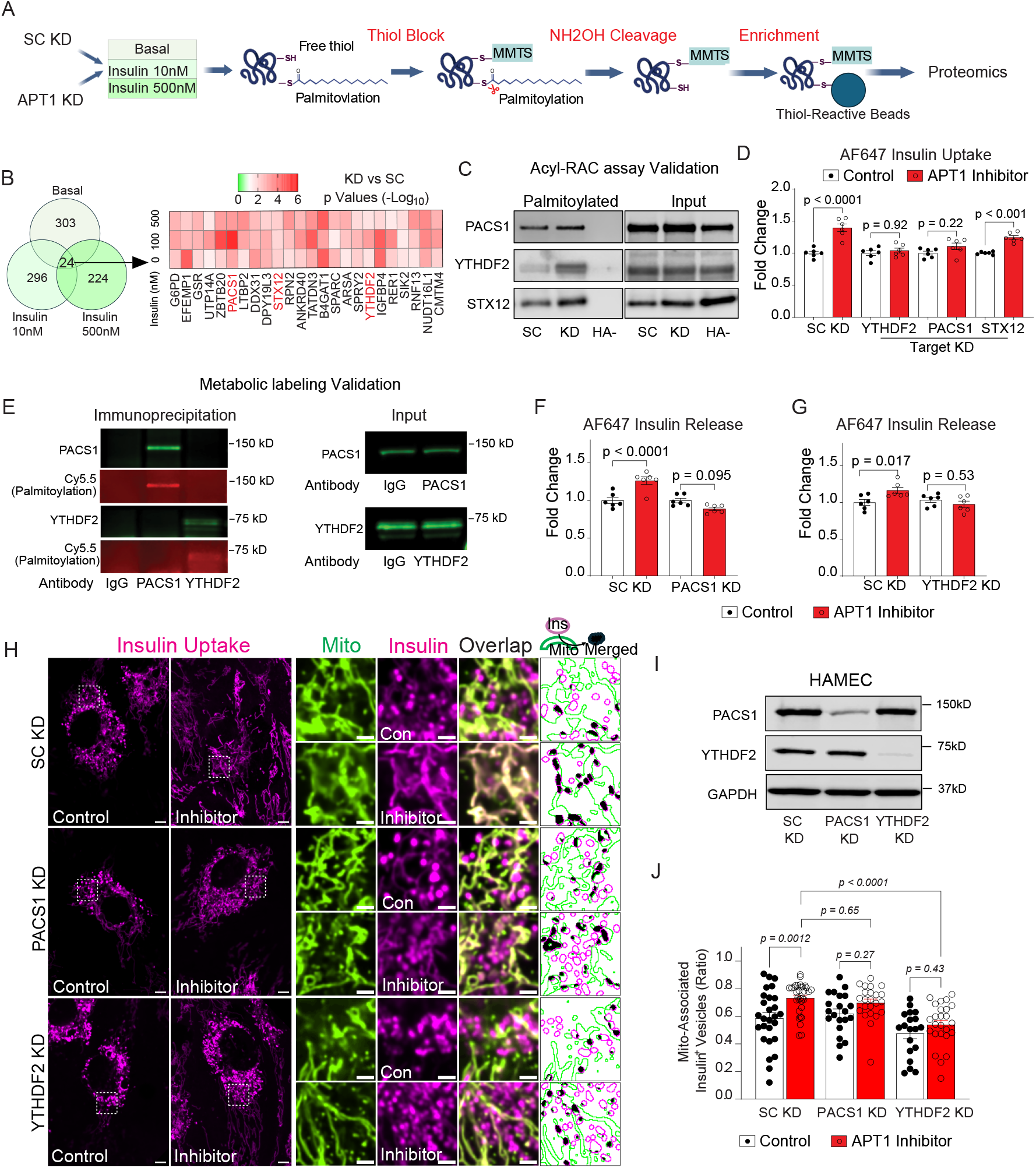
Palmitoylation proteomics identifies PACS1 and YTHDF2 as mediators of APT1-regulated insulin transport. (**A**) Schematic of palmitoylation proteomics workflow, showing acyl resin-assisted capture (RAC) enrichment prior to MS analysis. (**B**) Venn diagram showing the numbers of potential APT1 substrates identified in basal, 10 nM insulin, and 500 nM insulin conditions. 24 proteins were common to all 3 experiment groups (left), with heatmaps displaying the −log10(*p*) values for the comparison of KD versus SC among upregulated proteins under these conditions. (**C**) Validation of APT1-dependent PACS1, YTHDF2, and Syntaxin12 palmitoylation in HAMECs using RAC assay. HA-: no hydroxylamine cleavage control. (**D**) Effect of knocking down PACS1, YTHDF2, or STX12 on AF647 insulin uptake in HAMECs treated with or without APT1 inhibitor. (**E**) Palmitoylation of endogenous PACS1 and YTHDF2 confirmed by metabolic labeling followed by click chemistry. (**F-G**) Effect pf PACS1 (F) or YTHDF2 (G) knockdown on AF647 insulin release in HAMECs treated with or without APT1 inhibitor. (**H**) Representative images showing that PACS1 or YTHDF2 knockdown in HAMECs abolished APT1 inhibitor-induced association of vesicular AF647 insulin with mitochondria, under the same conditions as panels F and G. Scale = 5 μm (2 μm for zoomed-in images). (**I**) Knockdown of PACS1 and YTHDF2 in HAMECs confirmed by Western blot. (**J**) Quantification of mitochondria-associated insulin^+^ vesicles. Data are expressed as mean ± SEM. Statistical analyses were performed by two-way ANOVA with Sidak’s post hoc test. See also Figure S7.

## Discussion

Homeostasis requires the regulated movement of signaling molecules like insulin from a first space compartment, inside blood vessels, to the second space compartment of the interstitium surrounding cells. How insulin accesses metabolic tissues such as muscle and fat is not fully understood. Our results suggest that palmitoylation fine-tunes insulin transport across the capillary endothelium. Endothelial cell-specific deficiency of the depalmitoylase APT1 in mice ameliorates diet-induced glucose intolerance and insulin resistance by improving insulin delivery to the interstitium. Inhibition of APT1 in endothelial cells enhances insulin uptake independent of canonical endocytic pathways, with insulin translocating directly into mitochondria without detectable involvement of vesicular carriers. Subsequent transfer of insulin from mitochondria to endolysosomal compartments is APT1-dependent in part through effects on the APT1 targets YTHDF2 and PACS1. These findings suggest that insulin mitochondrial transport is involved in insulin sensitivity.

Capillary endothelial insulin transport can limit insulin-stimulated glucose disposal. A spatial and temporal gap between plasma and interstitial insulin levels has been documented in both animals and humans using lymph sampling,^8,12^ microdialysis,^11,17^ and intravital microscopy.^14,20^ This gap impacts the insulin response since interstitial insulin acts instantaneously on muscle cells to stimulate glucose uptake.^51^ The mitochondrial network, capable of exchange with the extracellular environment,^23,25^ enables endothelial cells to adapt to changes in circulating insulin from physiological peaks to pathological or pharmacological conditions. Through this buffering capacity, mitochondrial transport of insulin by endothelium may balance glucose regulation while preventing excessive insulin exposure.

Conflicting data address whether the insulin receptor is required for insulin transport. Radio-labeled iodinated insulin requires the insulin receptor for uptake by endothelial cells in vitro.^52^ In contrast, fluorescent-labeled insulin in mouse capillaries is transported by a non-saturable fluid-phase process that does not require the insulin receptor,^20^ confirming studies in dogs measuring insulin in hindlimb lymph derived from interstitial fluid surrounding muscle.^19^ Mixed results were seen in mice with endothelial insulin receptor knockouts.^17,18,53^ Part of the confusion may lie in the challenge of distinguishing changes in insulin transport from changes in perfusion, which is regulated by insulin receptors.^10,13,15,16,54^ The insulin receptor is minimally expressed in muscle capillaries, arguing against a receptor-dependent mechanism as the principal route for insulin transport.^16^ Even in brain with abundant capillary insulin receptors, insulin transport is largely unaffected by the lack of receptor-mediated signaling.^55^ We identified an APT1-dependent pathway for endothelial insulin transport with fluorescent-labeled insulin used to characterize fluid-phase transport in vivo^20^ and like fluid-phase transport, this pathway appears to be unsaturable and independent of the insulin receptor. Our data do not exclude other transport mechanisms at lower insulin concentrations. Endothelial insulin uptake pathways may differ depending on the insulin concentration.^56^ Our study supports an unsaturable insulin transport model allowing increased insulin delivery during pathological states.

Unlike previous models demonstrating insulin uptake by clathrin-coated pits or caveolae,^16,21,47,57^ our work suggests a specialized uptake in cultured endothelial cells that resembles, but is distinct from, pinocytosis. Live cell imaging showed labeled insulin immediately entering tubular networks co-localized with mitochondrial markers, and uptake was confirmed biochemically with native insulin. Mitochondria signal palmitoylation-dependent massive endocytosis,^29,30^ and proteins as well as large complexes such as lipoprotein particles internalized from the extracellular space can be transferred into mitochondria from endosomes,^28,58^ but we did not detect early endosomal involvement in insulin transport. This non-endosomal mediated uptake shares similarities with the direct membrane translocation observed for certain cell-penetrating peptides,^59^ for which the mechanisms of cellular entry are elusive. Given its sensitivity to Dyngo-4a, a potent dynamin inhibitor, mitochondrial uptake of insulin may involve regulators of mitochondrial membrane remodeling and tethering, such as Drp1 and Miro1. There are also precedents for mitochondria interacting with the extracellular space through the plasma membrane,^25^ consistent with our EM and TIRF imaging of endothelial mitochondria. Regulated positioning of mitochondria near the plasma membrane participates in membrane disruption.^60^ It is unclear if mitochondrial insulin uptake could be mediated through direct channeling from the plasma membrane. It is also unclear how mitochondrial insulin is transported out of cells. Potential mechanisms include mitochondria-derived vesicles,^61^ extracellular vesicles,^62^ secretory lysosomes,^63^ and membrane contact with or without direct release of mitochondria.^64^

APT1 affects mitochondrial biology, partially localizes to mitochondria,^39^ and in liver cells, coordinates the mitochondrial protein network for metabolic regulation,^40^ suggesting that palmitoylation/depalmitoylation cycling governs mitochondrial function. Differing from a previous report,^39^ the lack of APT1 enrichment in mitochondrial fractions of endothelial cells in the current study also points to a role for APT1 in a previously unrecognized function of mitochondria, insulin transport. Specifically, APT1 activity permits insulin transfer from mitochondria to late endosomes, and APT1 inhibition impairs this transfer, retaining insulin within mitochondria, which is correlated with enhanced insulin transport. These findings suggest that mitochondrial trafficking of insulin is essential for its transport. Consistent with this notion, cyclosporin A, known to protect mitochondrial integrity by inhibiting the mitochondrial permeability transition pore,^50^ preserves insulin content in mitochondria and enhances insulin transport in mice.

These data align with a prior report demonstrating that acute CsA administration in healthy humans enhances insulin sensitivity without affecting insulin secretion.^65^ In contrast, chronic CsA use as immunosuppressive therapy in transplant patients is associated with an increased risk of diabetes.^66^ These findings suggest that mitochondria-targeting actions of CsA may facilitate insulin transport, and raise the possibility that mitochondrial quality is a determinant of cellular insulin content. Our findings reveal a mitochondria-to-late-endosome trafficking axis for insulin in endothelial cells, likely mediated by direct organelle membrane contact or mitochondria-derived vesicles. This pathway appears to limit insulin transport and may be related to mitochondrial recycling. Endosome-mitochondria interactions participate in the clearance of damaged or dysfunctional mitochondria^23,67^ and Rab7 promotes mitochondria-lysosome contacts that support dynamic remodeling of the mitochondrial network.^68^

We identified two palmitoylated proteins required for APT1-mediated insulin transport. PACS1, a protein trafficking modulator, regulates endomembrane protein movement, mutations in its gene cause a neurodevelopmental syndrome,^69^ and it regulates the mitochondrial cell death pathway through mitochondrial outer membrane permeabilization.^70^ YTHDF2, a reader of m^6^-methyladenosine (m^6^A)- modified mRNAs, determines mRNA fate,^71^ and plays diverse roles in mitochondria including modulating membrane potential,^72^ regulating mitochondrial biogenesis,^73^ and promoting mitochondrial oxidative phosphorylation through m^6^A-independent mechanisms.^74^ YTHDF3, a paralog of YTHDF2, undergoes palmitoylation in its conserved YTH domain.^75^ Single nucleotide polymorphisms (SNPs) in the PACS1 gene are associated with severe obesity^76^ and YTHDF2 variants are associated with HbA1c^77^ and type 2 diabetes.^78^

In short, the endothelium maintains a palmitoylation-regulated insulin transport pathway through mitochondria. A better understanding of this pathway may yield novel therapeutics for insulin resistance and its associated metabolic disorders.

### Limitations of the study

In vitro experiments in this study utilized supraphysiologic insulin concentrations, an approach required by technical constraints that aligns with established experimental protocols for visualizing insulin transport. Several lines of evidence demonstrated that this palmitoylation-regulated endothelial mitochondrial insulin transport is a specific biological phenomenon rather than an artifact of high insulin concentrations. First, dose-response imaging revealed distinct mitochondrial insulin signals between 0.5 and 5 nM, a range spanning physiological postprandial states to severe insulin resistance. Second, both BID-induced mitochondrial insulin release and super-resolution microscopy excluded the possibility of non-specific mitochondrial binding. Third, pharmacological targeting of mitochondrial function via cyclosporin A in vivo increased endothelial insulin transport, elevating interstitial fluid insulin concentrations. Collectively, these data are consistent with physiological flexibility of the endothelium in modulating insulin transport across a broad range of exogenous insulin concentrations.

While this study identifies mitochondria-mediated transendothelial insulin transport, several components of this pathway are unclear. Specifically, the transport proteins or structural domains mediating insulin import into mitochondria, how mitochondria interact with the plasma membrane, and the details of insulin release into the interstitial space are unknown. Determining whether this process utilizes mitochondrial translocases, mitochondrial-derived vesicles, or inter-organelle contact sites could provide insight into endothelial transport relevant to insulin resistance.

## Resource availability

### Lead contact

Further information and requests for resources and reagents should be directed to and will be fulfilled by the lead contact, Clay F. Semenkovich.

### Materials availability

Plasmids generated in this study are available from the lead contact upon request.

### Data and code availability

All data reported in this paper will be shared by the lead contact upon request. The mass spectrometry proteomics data have been deposited to the ProteomeXchange Consortium via the PRIDE partner repository with the dataset identifier PXD074478. Western blot images for the figures in the manuscript are available as Data S1: Sources, Related to Figures 1-7, and S1–S7.

This paper does not report original code.

Any additional information required to analyze the data reported in this paper is available from the lead contact upon reasonable request.

## Acknowledgements

This work was supported by NIH grants HL157154, DK020579, DK056341, DK133344, UL1TR000448, the National Science Foundation (CBET-2316285 and CBET-2331330), the WashU Centene Personalized Medicine Initiative, the Irene and Michael Karl Professorship, and the Charles Kilo Diabetes Research Fund. We thank Chu Feng, Li Yin, and Petra Erdmann-Gilmore for expert technical support.

## Author contributions

W.Z. designed and conducted experiments, interpreted data, and wrote the manuscript. S.A., G.M., R.X., G.D., A.D., and Y.W. conducted experiments, interpreted data, and contributed to discussions. Q.Z. analyzed proteomics data. S.S. interpreted data and contributed to discussions. X.W. designed and conducted experiments, interpreted data, performed key analyses, and wrote the manuscript. C.F.S. designed experiments, obtained approvals, interpreted data, and wrote the manuscript. All authors had input into the manuscript.

## STAR★Methods

### Key resources table

| REAGENT or RESOURCE | SOURCE | IDENTIFIER |
| --- | --- | --- |
| **Antibodies** |  |  |
| Mouse monoclonal anti TOMM20 | Abcam | Cat# ab56783; RRID: AB_945896 |
| Rabbit polyclonal anti COX IV | Abcam | Cat# ab16056; RRID: AB_443304 |
| Rabbit monoclonal anti Lypla1 (APT1) | Abcam | Cat# ab91606; RRID: AB_10565192 |
| Mouse monoclonal anti total OXPHOS rodent cocktail | Abcam | Cat# ab110413; RRID: AB_2629281 |
| Rabbit polyclonal anti Syntaxin 12 | Proteintech | Cat# 14259-1-AP; RRID: AB_2198222 |
| Rabbit polyclonal anti phospho-AKT (Ser473) | Cell Signaling Technology | Cat# 9271; RRID: AB_329825 |
| Mouse monoclonal anti pan-AKT (40D4) | Cell Signaling Technology | Cat# 2920; RRID: AB_1147620 |
| Rabbit monoclonal anti Clathrin Heavy Chain (D3C6) | Cell Signaling Technology | Cat# 4796; RRID: AB_10828486 |
| Rabbit monoclonal anti Caveolin-1 (D46G3) | Cell Signaling Technology | Cat# 3267; RRID: AB_2275453 |
| Rabbit monoclonal anti GM130 (D6B1) | Cell Signaling Technology | Cat# 12480; RRID: AB_2797933 |
| Rabbit polyclonal anti β-tubulin | Cell Signaling Technology | Cat# 2146; RRID: AB_2210545 |
| Rabbit polyclonal anti α-Tubulin | Cell Signaling Technology | Cat# 2144; RRID: AB_2210548 |
| Rabbit monoclonal anti Cytochrome c (D18C7) | Cell Signaling Technology | Cat# 11940; RRID: AB_2637071 |
| Rabbit monoclonal anti YTHDF2 (E2I2H) | Cell Signaling Technology | Cat# 71283; RRID: AB_3068618 |
| Rabbit monoclonal anti Calnexin (C5C9) | Cell Signaling Technology | Cat# 2679; RRID: AB_2228381 |
| Rabbit polyclonal anti PACS1 | Atlas Antibodies | Cat# HPA038914; RRID: AB_10672831 |
| Mouse monoclonal anti GAPDH | Sigma | Cat# G8795; RRID: AB_1078991 |
| Goat polyclonal anti Mouse IgG (IRDye 680RD) | LicorBio | Cat# 926-68070; RRID: AB_10956588 |
| Goat polyclonal anti Rabbit IgG (IRDye 800CW) | LicorBio | Cat# 926-32211; RRID: AB_621843 |
| Rat monoclonal anti-mouse CD16/32 | BD Biosciences | Cat# 553142; RRID: AB_394657, |
| Rat monoclonal anti-mouse CD31 (APC) | BD Biosciences | Cat# 551262; RRID: AB_398497, |
| Rat monoclonal anti-mouse CD11b (APC/Cy7) | BioLegend | Cat# 101226; RRID: AB_830642, |
| Rat monoclonal anti-mouse F4/80 (PE) | Thermo Fisher Scientific | Cat# 12-4801-82; RRID: AB_465922 |
| Rat monoclonal anti-mouse CD45 (Pacific Orange) | Thermo Fisher Scientific | Cat# MCD4530; RRID: AB_2536417 |
| Anti-Insulin receptor $\alpha$, Clone47-9 | Leinco | Cat# 50103 |
| Anti-Human CD221/IGF1R, Clone 1H7 | BD Pharmingen | Cat# 555998 |
| **Bacterial and virus strains** |  |  |
| DH10B Competent Cells | Thermo Fisher Scientific | Cat# EC0113 |
| **Chemicals, peptides, and recombinant proteins** |  |  |
| S-methyl methanethiosulfonate (MMTS) | Sigma | Cat# 64306 |
| Hexadecylsulfonyl fluoride (HDSF) | Santa Cruz Biotechnology | Cat# sc-221708 |
| Hydroxylamine (HA) | Thermo Fisher Scientific | Cat# B22202.AE |
| Palmitic acid alkyne | Cayman Chemical | Cat# 13266 |
| Hexadimethrine bromide (Polybrene) | Sigma | Cat# H9268-10G |
| Tamoxifen | Sigma | Cat# T5648 |
| ML348 (APT1 inhibitor) | Tocris | Cat# 5345 |
| Humulin R U-100 | Eli Lilly and Company | Cat# 00002-8215-01 |
| Insulin Alexa fluor 647 labeled | Nanocs | Cat# IS1-AF647-1 |
| Insulin Cy5 labeled | Nanocs | Cat# IS1-S5-1 |
| Insulin, Human Recombinant (native) | Sigma | Cat# 91077C |
| Glucagon (1-29)-[Lys(AF647)] | Biosynth | Cat# CRB1101573 |
| Recombinant Human BID (Caspase-8-cleaved) | R&D systems | Cat# 882-B8 |
| Human IGF1 | Thermo Fisher Scientific | Cat# AF100-11-100UG |
| **Critical commercial assays** |  |  |
| Human Insulin Antibody Pair - BSA and Azide free | Abcam | Cat# ab253508 |
| Human Insulin ELISA kit | Alpco | Cat# INSHU80-INSHU-E01.1 |
| PureLink RNA Mini Kit (Thermo) | Thermo Fisher Scientific | Cat# 12183018A |
| iScript cDNA synthesis kit | Bio-Rad | Cat# 1708890 |
| SYBR Premix Ex Taq | TAKARA BIO | Cat# RR420A |
| Lenti-X concentrator | TAKARA BIO | Cat# 631231 |
| BCA protein assay kits | Thermo Fisher Scientific | Cat# 23227 |
| **Deposited data** |  |  |
| Mass spectrometry dataset | ProteomeXchange | Accession ID: PXD074478 |
| **Experimental models: Cell lines** |  |  |
| Human Adipose Microvascular Endothelial Cells (HAMECs) | ScienCell | Cat# 7200 |
| HEK293T cells | ATCC | Cat# CRL-11268 |
| **Experimental models: Organisms/strains** |  |  |
| Mouse: C57BL/6J floxed Lypla1 | Wei et al.<sup>36</sup> | N/A |
| Mouse: C57BL/6J floxed Lypla2 | Wei et al.<sup>40</sup> | N/A |
| Mouse: Tie2-cre mice | Jackson Laboratory | B6.Cg-Tg(Tek-Cre)12Flv |
| Mouse: VE-cadherin-Cre-ERT2 mice | Ralf H Adams lab | Tg(Cdh5-cre/ERT2)1Rha |
| Mouse: LysM-Cre mice | Jackson Laboratory | B6.129P2-Lyz2tm1(cre)lfo/J |
| **Oligonucleotides** |  |  |
| Primers used for qPCR | Table S1 | N/A |
| siRNA sequences | Table S2 | N/A |
| **Recombinant DNA** |  |  |
| pLVX-EGFP-Rab11b | Addgene | 134860 |
| pLVX-EGFP-Rab5a | Addgene | 134858 |
| pLVX-EGFP-Rab7a | Addgene | 133027 |
| pLVX-EGFP-Lamp1 | Addgene | 134868 |
| pLVX-EGFP-COX8 | Addgene | 134861 |
| pLVX-EGFP-MAO | Addgene | 133025 |
| pLV-mScarlet-I | Addgene | 159172 |
| psPAX2 | Addgene | 12260 |
| pMD2.G | Addgene | 12259 |
| **Software and algorithms** |  |  |
| Mascot software10 version 2.8.0.1 | Perkins et al. | http://www.matrixscience.com/ |
| R | R Core Team (v4.3.0, 2023) | https://www.R-project.org/ |
| Posit/RStudio | RStudio Team (2023.03.0) | https://posit.co/products/open-source/rstudio/ |
| R package: limma | Ritchie et al.<sup>79</sup> | bioconductor.org |
| R package: mixtools: normalmixEM | Benaglia et al. | http://www.jstatsoft.org/v32/i06/ |
| proteoQ | Zhang Q (v1.7.5.1, 2023) | https://github.com/qzhang503/proteoQ |
| FlowJo_V10.8 | Flowjo LLC. | https://www.flowjo.com/ |
| Image Studio Lite Ver 5.2 | Licor | https://www.licor.com/bio/image-studio/ |
| Prism 11 | Graphpad software | https://www.graphpad.com/scientific-software/prism/ |
| Fiji-ImageJ | National Inst. Of Health | https://imagej.nih.gov/ij/ |
| Fiji plugin: Trackmate | Tinevez et al.<sup>80</sup> | https://imagej.net/TrackMate |
| **Other** |  |  |
| Lectin-Dylight649 | Vector labs | Cat# DL-1178-1 |
| Texas Red-dextran 70kDa | Thermo Fisher Scientific | Cat# D1830 |
| Texas Red-dextran 3kDa | Thermo Fisher Scientific | Cat# D3329 |
| Dil-LDL | Thermo Fisher Scientific | Cat# L3482 |
| AF647 transferrin | Thermo Fisher Scientific | Cat# T23366 |
| Cy5.5 Azide | Vector labs | Cat# CCT-1059-1 |
| Fibronectin | Sigma | Cat# F1141 |
| Lipofectamine 3000 | Thermo Fisher Scientific | Cat# L3000008 |
| Live Cell Imaging Solution | Invitrogen | Cat# A59688DJ |
| EGM-2 MV media | Lonza | Cat# CC-3202 |
| DMEM | Thermo Fisher Scientific | Cat# 11965092 |
| High fat diet | Envigo | Cat# TD88137 |
| Intercept blocking buffer | Licor | Cat# 927-60001 |
| TOM22 magnetic beads | Miltenyi Biotec | Cat# 130-094-833 |
| Thiopropyl sepharose beads | Nanocs | Cat# AR-S3-1 |
| Magnetic protein G Dynabeads | Thermo Fisher Scientific | Cat# 10007D |
| Contour glucose strips | Bayer | Cat# 7080G |
| DharmaFECT 1 Transfection Reagent | Horizon Discovery | Cat# T-2001-02 |
| Novex Wedge well 4-20% Tris-glycine mini gels | Thermo | Cat# XP04200BOX |
| PVDF membrane (0.45 µm) | Millipore | Cat# IPFL00010 |
| TransIt-LT1 transfection reagent | Mirus | Cat# MIR 2300 |
| Cyclosporin A | Sigma | Cat# SML1018 |
| MitoTracker™ Green FM | Thermo Fisher Scientific | Cat# M7514 |
| cOmplete™, EDTA-free Protease Inhibitor Cocktail | Roche | Cat# 11873580001 |
| PhosSTOP™ phosphatase inhibitor tablets | Roche | Cat# 4906845001 |
| DAPI (Viability Dye) | BD Biosciences | Cat# 564907 |

### Experimental Model and Study Participant Details

#### Animal models

Animal protocols were approved by the Institutional Animal Care and Use Committee (IACUC) at Washington University in St. Louis and followed the NIH Guide for the Care and Use of Laboratory Animals. Mice floxed for *Lypla1* or *Lypla2* (the genes for APT1 and APT2, respectively) were generated on a C57BL/6J background as described.^36^ Mice with endothelial cell-specific knockout of APT1 or APT2 (Tie2-APT1/2 KO and VEC-APT1/2 KO) were generated by crossing floxed mice with constitutive Tie2-Cre mice (B6.Cg-Tg(Tek-Cre)12Flv), or VE-cadherin-Cre-ERT2 mice (Tg(Cdh5-Cre/ERT2)1Rha). Inducible deletion was achieved with tamoxifen (2 mg per mouse IP) once daily for five consecutive days starting at 6 weeks of age. Mice with macrophage-specific knockout of APT1 were generated with LysM-Cre mice (B6.129P2-Lyz2tm1(Cre)Ifo/J). Animals resided in a specific pathogen-free facility with a 12h light-dark cycle and were fed standard chow diet or a high-fat diet with 42% calories from fat (Envigo, TD.88137). Knockout and littermate control mice aged 4-9 months were studied. Both male and female mice were included in the study. Only male data are presented, as the phenotype of improved insulin sensitivity in knockout mice was observed exclusively in males, consistent with prior reports that male mice are more susceptible to HFD-induced metabolic dysfunction.^81^

#### Cell lines

HAMECs (Human Adipose Microvascular Endothelial Cells) were obtained from ScienCell, grown in EGM-2 MV media (Lonza, CC-3202, containing 5 mM D-glucose) and used at passage number <10. Culture plates were coated with fibronectin (Sigma, F1141) before use. Human HEK293T cells were obtained from ATCC (Catalog No. CRL-11268) and maintained in high-glucose DMEM containing 10% FBS. Cells were cultured at 37°C and 5% CO2 and tested negative for mycoplasma.

### Method Details

#### Metabolic phenotyping

Body composition was determined using Echo-MRI. For fasting plasma collection, glucose (GTT), or insulin tolerance tests (ITT), animals were fasted for 4-6h.^82^ Mice were injected IP with 1 g/kg glucose (GTT) or 0.75 IU/kg insulin (ITT). Blood glucose was measured at 0, 15, 30, 60, and 120 min post injection with a Contour glucose meter (Bayer). To measure insulin-stimulated AKT signaling in tissues,^82^ mice were fasted overnight, weighed, and then anesthetized with a ketamine/xylazine cocktail. Anesthetized mice were injected with 5 IU/kg insulin, and 10 min later were perfused with cold PBS followed by tissue collection.

#### Flow cytometric analysis for adipose tissue-associated macrophages

Epididymal fat pads were isolated after saline perfusion, then minced, washed, and centrifuged (1,000 g for 5 min). Floating adipose tissue was collected and digested with type I collagenase solution (Gibco, 1 mg/ml, 30 min). This digest was pelleted to yield the stromal vascular fraction (SVF), which was washed twice in FACS wash buffer (PBS supplemented with 4% FBS), then blocked for Fc receptors using anti-mouse CD16/32 (clone 2.4G2, BD Bioscience), followed by staining with APC-conjugated anti-mouse CD31 (MEC 13.3, BD Bioscience), APC/Cy7-conjugated anti-mouse CD11b (clone M1/70, Biolegend), PE-conjugated anti-mouse F4/80 (clone BM8, Thermo Fisher Scientific), Pacific Orange-conjugated anti-mouse CD45 (clone 30-F11, Thermo Fisher Scientific), and DAPI (BD Bioscience). The emission fluorescence of the cell suspension was acquired by digital FACScan (Cytek Development), and data analysis was performed in FlowJo.

#### Quantitative PCR

Tissues were isolated from mice perfused with cold PBS followed by snap-freezing in liquid nitrogen and storage at –80°C. RNA was isolated using PureLink RNA Mini Kit (Thermo 12183018A) with on-column gDNA digestion (Thermo 12185010). One microgram total RNA was used to synthesize cDNA using the iScript cDNA synthesis kit (Bio-Rad, Cat. NO.1708890). Gene expression was measured with Takara Green Premix Ex Taq (RR420A) in a QuantStudio 3 System (Applied Biosystems). TBP was used as reference for normalization. The primers for mouse tissue are listed in Table S1.

#### Western blotting

Protein samples were loaded for SDS-PAGE (Novex Wedge well 4-20% Tris-glycine mini gels, Thermo XP04200BOX) and run at 120-140V for 90 min. After wet transfer to PVDF (0.45 µm, Millipore, IPFL00010), membranes were blocked (Licor 927-60001) for 60 min at RT. 0.1% Tween-20 and primary antibodies were added to membranes and incubated overnight at 4°C. Secondary antibodies were incubated with membranes in 50% Licor Odyssey blocking buffer with 0.1% Tween-20 at RT for 60 min. Membranes were imaged with a Licor Odyssey Fc imaging system, and protein intensity was quantified with Licor Image Studio software. The following antibodies were used with dilution 1:1000-1:2000: TOMM20, Abcam, ab56783; COX IV, Abcam, ab16056; Lypla1 (APT1), Abcam, ab91606; total OXPHOS rodent antibody cocktail, Abcam, ab110413; Syntaxin 12, Proteintech, 14259-1-AP; phospho-AKT (Ser473), Cell Signaling Technology, 9271; pan-AKT (40D4), Cell Signaling Technology, 2920; Clathrin Heavy Chain (D3C6), Cell Signaling Technology, 4796; Caveolin-1 (D46G3), Cell Signaling Technology, 3267; GM130 (D6B1), Cell Signaling Technology, 12480; β-tubulin, Cell Signaling Technology, 2146; α-Tubulin, Cell Signaling Technology, 2144; Cytochrome c (D18C7), Cell Signaling Technology, 11940; YTHDF2 (E2I2H), Cell Signaling Technology, 71283; Calnexin (C5C9), Cell Signaling Technology, 2679; PACS1, Atlas Antibodies, HPA038914; GAPDH, Sigma, G8795; IRDye 680RD Goat anti-Mouse IgG secondary antibody, LicorBio, 926-68070; IRDye 800CW Goat anti-Rabbit IgG secondary antibody, LicorBio, 926-32211.

#### Fluorescence labeled insulin bioactivity confirmation

Fluorescence labeled insulins (AF647 insulin and Cy5 insulin) were tested for bioactivity by AKT phosphorylation in HAMECs. HAMECs were serum starved for 3 h in 1% BSA/EBM-2 media, treated with various doses of insulin for 15 min, and cell lysates were prepared. Phosphorylation of AKT was measured by Western blot. Unlabeled native human insulin was used as positive control.

#### Vessel density

For histological staining of skeletal muscle vasculature, mice under general anesthesia were sequentially perfused through the heart with heparinized normal saline (10 U/ml heparin), a vasodilation solution (10 μM sodium nitroprusside), and 4% PFA (paraformaldehyde in PBS). Lower calf muscle tissues were then collected and fixed in 4% PFA (overnight at 4 °C) followed by cryoprotection with 30% sucrose (48 h at 4 °C). Frozen sections were prepared using a Leica CM1850 Cryostat and stained with lectin-Dylight649 (Vector labs). Images were acquired using an automated fluorescence slide scaner (Zeiss AxioScan 7), and numbers/areas of blood vessels were analyzed in QuPath.

#### Gene knockdown

For siRNA experiments in HAMECs, DharmaFECT 1 Transfection Reagent (2 μl/mL, Horizon Discovery, T-2001-02) was used. Synthesized siRNA oligonucleotides were as follows: non-targeting siRNA control (NTC, Horizon Discovery, D-001810-10-20) or ON-TARGETplus SMARTpool siRNA (Horizon Discovery) for APT1 (L-010007-00-0005), PACS1(L-006697-01-0005), YTHDF2 (L-021009-02-0005), and STX12 (L-018246-01-0005). ON-TARGETplus individual siRNAs were obtained for Clathrin HC (J-004001-09-0002) and Caveolin 1 (J-003467-06-0002). siRNA concentration was 10-20 nM. Cells were studied 72-96 h after transfection.

#### Plasmids

Plasmids used in imaging were obtained from Addgene: pLVX-EGFP-Rab11b (134860), pLVX-EGFP-Rab5a (134858), pLVX-EGFP-Rab7a (133027), pLVX-EGFP-Lamp1 (134868), pLVX-EGFP-COX8 (134861), and pLVX-EGFP-MAO (133025). An mScarlet-conjugated APT1 expression construct was derived from pLV-mScarlet-I (Addgene, 159172). Lentivirus was generated in 293T cells by transfection of vectors with packaging helper plasmids psPAX2 (Addgene, 12260) and pMD2.G (Addgene, 12259). HAMECs were infected for 24 h with virus collected from filtered supernatants of 293T cells 48 h after transfection in the presence of Polybrene (10 µg/ml).

#### Fluorescence live-cell imaging and image analysis

Live-cell images were acquired by a Nikon spinning disk confocal system (Plan Apo λ 60x oil/NA 1.4, LU-NV NIDAQ MultilLaser, Yokagawa CSU-X1 Nipkow spinning disk scan, Fusion sCMOS camera, ultra-quiet mode). Cells were cultured in glass bottomed chambers and changed to live cell imaging solution (Thermo) before imaging by a Ti-E microscope equipped with a Tokai-hit stage-top incubator capable of temperature and humidity regulation. Fluorescent-labeled insulin or other endocytic markers were loaded into cells for 10 to 15 min at 37 °C, followed by 2 washes with imaging buffer. The perfect focus stabilization (PFS) system was engaged during imaging acquisition. Images were then deconvoluted with the Denoise.ai module in NIS-Elements and subsequently transferred to Fiji for visualization and analysis. For PFA fixation, cells were fixed in 1% PFA in PBS at 37 °C for 10 min. For cold-start experiments, cells were kept on ice during the insulin-loading phase, washed with cold imaging buffer, and maintained at low temperature in the imaging chamber using an ice-water mixture before temperature elevation. For APT1 inhibition, cells were pre-treated with 10 μM ML348 for 2-4 h. Live-cell TIRF images were acquired by a Nikon TIRF system equipped with the PerfectFocus stability mechanism (SR Apo TIRF 100x oil/NA1.49, Andor Zyla camera 4.2 Megapixel sCMOS camera VSC-03228, Ti-E microscope, LU-NV NIDAQ STORM QUAD).

Structured illumination microscopy (SIM) super-resolution images were taken on a Nikon N-SIM microscope equipped with a 100× objective lens (1.49 NA, Nikon). Images were captured using Nikon NIS-Elements and reconstructed using slice reconstruction in NIS-elements. All Images were taken at a single Z-plane. Cells were cultured in live cell imaging buffer and maintained in a temperature-controlled chamber (37 °C) at 5% CO2 in a TokaiHit stage top incubator.

To analyze insulin vesicle movement, image series were analyzed in Fiji by TrackMate plugin ^80^. Randomly selected regions (250 x 250 pixels, 100-second video) adjacent to cell nuclei were analyzed with the same particle-detection settings (diameter = 0.5 and threshold = 5) and tracking configuration (simple LAP tracker model with linking max distance = 0.5 and no gap). Tracks with short duration (< 3 sec) were filtered out, and results from individual cells were averaged and recorded.

To analyze mitochondria-associated insulin vesicles, images captured at 37 °C were processed in Fiji. Mitochondrial areas labeled by GFP-conjugated markers were segmented using auto-thresholding. Insulin vesicles were segmented using FeatureJ Laplacian filter (smoothing = 1) followed by auto-thresholding. The ratio of vesicle area overlapping with mitochondria relative to total vesicle area was calculated and reported.

#### Insulin uptake and release assay

HAMECs were cultured on fibronectin-coated 24-well plates to confluence in complete EGM-2 MV media, 500 nM native human insulin (Sigma, I0908) or fluorescent-labeled insulin (Alexa fluor 647 labeled Insulin, Nanocs, IS1-AF647-1) was added, then after 15 min cells were washed with PBS twice, and cell lysates were measured by ELISA (Alpco, 80-INSHU-E01.1) for insulin or plate reader for fluorescence intensity. Insulin content was corrected by total protein with BCA assay (Thermo Fisher Scientific, 23227). In some experiments, cells were pre-treated with 10 μM APT1 inhibitor ML348 (Tocris, 5345) for 4 h. For competition experiments, 10x or 100x unlabeled native insulin, 100x IGF1 (Thermo Fisher Scientific, 100-11-100UG), or blocking antibodies (Anti-Insulin receptor α Clone 47-9, Leinco, 50103; Anti-Human CD221/IGF1R, Clone 1H7, BD Pharmingen, 555998) were added 5 min before the fluorescent-labeled insulin was loaded to HAMECs. For the insulin release assay, cells were loaded with insulin for 15 min, washed twice with PBS, new release buffer (EBM-2 basal media with 3% BSA, Sigma, A7030) was added and 15 min later, media were collected for insulin measurement^47^. The insulin receptor antagonist peptide S961 was kindly provided by Novo Nordisk.

#### Glucagon, LDL, transferrin and dextran uptake assay

For glucagon uptake, cells were treated with 500nM AF647 glucagon for 15 min in the presence or absence of APT1 inhibitor. Uptake of LDL, transferrin and dextran was performed as with the insulin uptake assay. HAMECs were loaded with Dil-LDL (5 μg/mL, Thermo Fisher, L3482), AF647 transferrin (25 μg/mL, Thermo Fisher, T23366), Texas Red-dextran 70kDa (25 μg/mL, Thermo Fisher, D1830), or Texas Red-dextran 3kDa (25 μg/mL, Thermo Fisher, D3329) for 15min.

#### Microneedle patch assay for ISF insulin

Quantification of insulin in interstitial fluid (ISF) was performed as described ^46^. Microneedle patches were coated with anti-Insulin capture antibody (2 μg/mL in PBS overnight, Human Insulin DuoSet kit, R&D systems, DY8056-05), followed by washing with PBST and blocking (3% BSA in PBS) for 1 h. To measure insulin in dermal ISF, mice were fasted for 4 h and anesthetized with isoflurane. Subsequently, mice were injected IP with 5 IU/kg human insulin (Eli Lilly, Humulin R U-100) then after 10 min microneedle patches were applied to mice skin for an additional 20 min. For the basal and 30 min after insulin conditions, mouse tail blood glucose was measured. Removed patches were blocked (3% BSA in PBS) for 1 h and incubated with biotinylated detection antibody, followed by streptavidin–800CW plasmonic fluor conjugate binding. Patch washes were finished in 24-well plate on a strong magnetic base. Fluorescence images of microneedle patches were obtained using a LI-COR fluorescence imager with the following scanning parameters: laser power 2-4; resolution 21 μm; channel: 800; height: 0 mm.

#### Skin ISF isolation

Mice were fasted for 4 h and IP injected with insulin at 5 IU/kg. After 30 min, mice were sacrificed and abdominal skin (∼1 inch square) was dissected. Skin samples were blotted gently, placed on 15-micron nylon mesh (ELKO filtering Co. 03-15/10), then tightly held by rubber band on a 50 mL conical centrifuge tube. ISF was isolated by centrifugation at 10000 x *g* for 20 min at 4°C as described ^83,84^. Insulin was measured by ELISA. To evaluate the effect of Cyclosporin A (CsA) on insulin transport, wild type male mice were injected IP once with 30 mg/kg CsA (in corn oil, Sigma, C2163000). Ten mins later, 2 IU/kg insulin was injected IP followed by isolation of skin ISF.

#### Transmission electron microscopy

Mice under anesthesia were sequentially perfused through the heart with heparinized normal saline (10 U/ml heparin), and 4% paraformaldehyde (PFA) with 0.1% glutaraldehyde (GA). The soleus muscle was carefully dissected without pulling or stretching and placed into 2 mL TEM fixative (2% PFA plus 2.5% GA, in 0.15 M cacodylate buffer, pH 7.2). Tissue fixation was continued overnight at 4 °C using a shaker. Samples were then rinsed in cacodylate buffer 3 times for 10 min each and subjected to a secondary fixation for one hour in 2% osmium tetroxide/1.5% potassium ferrocyanide in cacodylate buffer. Samples were rinsed in ultrapure water 3 times for 10 min and stained overnight in an aqueous solution of 1% uranyl acetate at 4°C. Then amples were washed in ultrapure water 3 times for 10 min each, dehydrated in a graded acetone series (50%, 70%, 90%, 100% x 3) for 10 min at each step, and infiltrated with microwave assistance (Pelco BioWave Pro) into LX112 epon resin. Samples were cured at 60°C for 72 h and ultrathin sections of 90 nm were cut, stained with uranyl acetate/lead citrate and imaged with a JEOL JEM-1400 Plus Transmission Electron Microscope using an AMT Nanosprint15-MkII sCMOS camera.

For muscle capillary endothelium vesicle quantification, two control and two Tie2 APT1 KO mice on HFD were used. For each mouse soleus sample, 3-4 images were quantified manually with ImageJ software. Total vesicle number, vesicle density, and specified abluminal vesicle number were determined. Capillary endothelium was identified based on a single endothelial cell layer, tight junctions, basement membranes, and intracellular vesicles.

#### Mitochondrial isolation

Mitochondria from HAMECs were isolated by centrifugation ^85^. HAMECs were lysed in mitochondria isolation buffer (MIB): 225mM mannitol, 75mM sucrose, 0.1 mM EGTA, and 30 mM Tris–HCl pH 7.4 and homogenized with a 25 G syringe for 10 strokes. Lysates were centrifuged at 600 x *g* for 5 min at 4°C, the pellet fraction was retained, and the supernatant was centrifuged at 7000 x *g* for 10 min at 4°C to generate a supernatant fraction. The pellet was washed with MIB then subjected to centrifugation at 10000 x *g* for 10 min at 4°C. The final mitochondrial fraction was used immediately or stored at −80°C. In some experiments, mitochondria were isolated using TOM22 magnetic beads (Miltenyi Biotec, 130-094-833).

#### BID induced insulin release from mitochondria

Recombinant Human BID (Caspase-8-cleaved, R&D systems, 882-B8) was used to induce mitochondrial outer membrane permeabilization (MOMP) with readout as cytochrome C release from mitochondria. To measure insulin release from mitochondria, HAMECs were treated with 500 nM native insulin or AF647 insulin for 15 min, and mitochondria were isolated as described above. Isolated mitochondria were diluted in MOMP buffer (125 mM KCl, 0.5 mM MgCl2, 3.0 mM succinic acid, 3.0 mM glutamic acid, 10 mM HEPES-KOH, 1 mg/mL BSA, proteinase inhibitor cocktail, pH 7.4) at 2 mg/mL. 75 μl mitochondria were used with or without 100 nM BID treatment at 30°C for 30 min. Vehicle control group represents the spontaneous cytochrome C release during incubation in MOMP buffer, and another control group consisted of mitochondria without incubation (NT, no treatment) to assess total cytochrome C content. After BID treatment, samples were centrifuged at 10000 x *g* for 10min, the supernatant was retained as release buffer fraction, and the mitochondrial pellet was lysed with RIPA buffer. Insulin in each fraction was measured by ELISA (for native insulin) or plate reader (for AF647 insulin). Data were corrected by total protein content.

#### ELISAs

Human insulin ELISA kits were purchased from Alpco (80-INSHU-E01.1). Manufacturer’s protocols were used for all plasma or media measurements.

#### Acyl-RAC assay

An acyl-RAC assay was performed to enrich palmitoylated protein as described ^36,37,40^. Cells were homogenized in lysis buffer (150 mM NaCl, 50 mM Tris, 5 mM EDTA, 2% Triton-X-100, 0.2 mM HDSF, pH 7.4, and protease inhibitor cocktail from Roche). Following sonication on ice, protein concentrations were determined by BCA protein assay kits (Thermo), and the same amounts of protein lysates were diluted to 2.5% SDS with 0.1% MMTS (S-Methyl methanethiosulfonate), and incubated at 37°C for 20 min. Proteins were then precipitated with ice-cold acetone (1:3 v/v protein solution: acetone) at −20°C for 30 min. Protein precipitates were obtained by centrifuging at 12000 rpm for 10 min, washed 3 times with cold 70% acetone, and air dried. Pellets were then resuspended with binding buffer (100mM HEPES, 1 mM EDTA, 1% SDS, pH=7.4, and protease inhibitor cocktail from Roche) by vortexing for 20 min at room temperature. Solubilized fractions were obtained by centrifuging at 14000rpm for 5 min, and supernatants were then aliquoted into 2 tubes containing thiopropyl sepharose beads (Nanocs, AR-S3-1), with or without freshly-prepared HA (hydroxylamine 0.8 M, pH=7.4), and rotated at room temperature for 2 h. Beads were washed 5 times with binding buffer, and enriched proteins were eluted with binding buffer containing 50mM DTT (20 min incubation at room temperature). Enriched palmitoylated proteins were subjected to proteomics or analyzed by Western blot using Licor infrared fluorescence imaging.

#### Palmitoyl-proteomics

HAMECs were cultured in 0.5% FBS EBM-2 media (Lonza, CC3156) for 4 h and treated with vehicle or insulin (10 or 500 nM, Sigma, I0908) for 15 min. Cell lysates were prepared and the palmitoylated proteins were enriched using acyl-RAC beads, then eluted protein samples were trypsin-digested and analyzed by label-free mass spectrometry ^36,41^.

Peptides were separated using a nano-ELUTE chromatograph (Bruker Daltonics, Bremen, Germany) and analyzed with trapped ion mobility time-of-flight mass spectrometry. Data from the mass spectrometer were converted to peak lists using DataAnalysis (version 5.2, Bruker Daltonics). MS2 spectra with parent ion charge states of +2, +3 and +4 were analyzed using Mascot software ^86^ (Matrix Science, version 2.8.0.1) against a concatenated UniProt (version January 2023) database of human and common contaminant proteins (cRAP, version 1.0 Jan. 1st, 2012; 116 entries). PSMs were filtered at 1% false-discovery rate (FDR) by searching against a reversed database. A minimum of two peptides with unique sequences, not resulting from missed cleavages, was required for identification of a protein. The processing, quality assurance and analysis of LC-MS data were performed with proteoQ (version 1.7.5.1, https://github.com/qzhang503/proteoQ). software developed with the tidyverse approach (https://CRAN.R-project.org/package=tidyverse) with open source software for statistical computing and graphics, R (https://www.R-project.org/) and RStudio (http://www.rstudio.com/). Metric multidimensional scaling (MDS) and principal component analysis (PCA) of protein log2-ratios were performed with the base R function stats:cmdscale and stats:prcomp, respectively. Heat-map visualization of protein log2-ratios was performed with heatmap. Linear modelings were performed using the contrast fit approach in limma ^79^, to assess statistical significance of protein abundance differences between indicated groups of contrasts.

#### Metabolic labeling and click chemistry

Protein palmitoylation was confirmed in HAMECs by metabolic labeling combined with click chemistry. Cells were incubated with media containing 20 μM palmitic acid alkyne (Cayman chemical, Cat. No. 13266) for 16 h, then lysed, and solubilized protein fractions were isolated. The same amounts of protein lysates were immunoprecipitated with 2 μg antibody-mediated magnetic protein G Dynabeads (Thermo Fisher Scientific, 10007D). Immunoprecipitates were click-labeled on beads with labeling buffer (PBS, pH 7.4, 1 mM CuSO4, 1 mM TCEP, 0.1 mM TBTA, 40 μM azide-Cy5.5, Vector labs) at room temperature for 1 h. Labeled proteins were blotted using primary antibodies and IRDye-800CW conjugated secondary antibody (Licor). Resultant blots were scanned by a dual-color fluorescence Licor imager.

#### Statistical analysis

All statistical analyses were performed in Prism 11 software (GraphPad Software) and p < 0.05 was considered significant. Comparison data are represented by scatter dot plot (with mean ± SEM) and analyzed by unpaired t test (with or without Welch’s correction) or two-way ANOVA with post hoc tests.

For animal studies, results are pooled data points, each corresponding to an individual mouse (n indicates mouse numbers). For histology experiments, each data point represents the mean value from randomly selected tissue sections of comparable areas for each mouse. For TEM experiments, each data point corresponds to an individual capillary endothelial cell analyzed from two pairs of mice. For statistical analysis, GTT and ITT data were analyzed by two-way ANOVA (matched values across time points) with Sidak’s test for multiple comparisons; body composition, adipose macrophage infiltration, gene expression, ISF insulin, AKT phosphorylation, muscle vessel quantification, and intracellular vesicle characterization were analyzed by unpaired t test with Welch’s correction.

For cell-based studies, results are pooled data points, with n (data points number in scatter dot plot) described as follows. For insulin uptake and release, endocytic marker uptake, mitochondria isolation and permeabilization assays, n refers to the number of replicates performed for each condition. For the quantification of mitochondria-associated insulin vesicles, each data point corresponds to an individual cell imaged across at least two independent experiments. For vesicle-movement analysis, Tukey plots represent the distribution of each track, and the mean values for each cell obtained from two independent experiments are shown in scatter plots. Most comparisons were performed by unpaired t test with Welch’s correction.

