## Supplements for "Palmitoylation calibrates mitochondrial transport of insulin across the endothelium in insulin resistance"

### Supplemental Information

### Supplemental Tables S1-S2

### Supplemental Figures S1-S7

### Video Descriptions

**Video S1. Related to Figure 5.**
Representative time-lapse confocal image series of HAMECs expressing the mitochondrial marker GFP-COX8 (green) and loaded with Alexa Fluor 647-labeled human insulin (magenta). Arrows highlight the rapid exit of insulin from mitochondria into peri-mitochondrial vesicles. Timestamps (mm:ss) indicate elapsed time after warming. Scale = 2 μm.

**Video S2. Related to Figure 5.**
Representative time-lapse confocal image series of HAMECs expressing the late endosome marker GFP-Rab7a (green) and loaded with Alexa Fluor 647-labeled human insulin (magenta). Arrows highlight insulin transfer from tubular structures into Rab7a-positive endosomes. Timestamps (mm:ss) indicate elapsed time after warming. Scale = 2 μm.

**Video S3. Related to Figure 6.**
Representative time-lapse confocal image series of Alexa Fluor 647-labeled human insulin-containing vesicles (gray) in HAMECs with control treatment (SC) or APT1 knockdown (KD), acquired at 37°C at 1 fps (frame per second). Vesicle trajectories are overlaid and color-coded by displacement to illustrate insulin vesicle mobility. Color scale: 0–3 μm displacement. Scale = 2 μm.

**Table S1. qPCR Primers used in the study.**

| Gene | Direction | Sequence |
|---|---|---|
| CD31 F | Forward | ACGCTGGTGCTCTATGCAAG |
| CD31 R | Reverse | TCAGTTGCTGCCCATTCATCA |
| VEGFR1 F | Forward | CCACCTCTCTATCCGCTGG |
| VEGFR1 R | Reverse | ACCAATGTGCTAACCGTCTTATT |
| IL12 F | Forward | CTCAGAAGCTAACCATCTCCTGG |
| IL12 R | Reverse | CACAGGTGAGGTTCACTGTTTC |
| IL1b F | Forward | GAAATGCCACCTTTTGACAGTG |
| IL1b R | Reverse | CTGGATGCTCTCATCAGGACA |
| IFNg F | Forward | ATGAACGCTACACACTGCATC |
| IFNg R | Reverse | TCTAGGCTTTCAATGACTGTGC |
| IL-17 F | Forward | TCAGCGTGTCCAAACACTGAG |
| IL17 R | Reverse | GACTTTGAGGTTGACCTTCACAT |
| IL-6 F | Forward | ATGGATGCTACCAAACTGGAT |
| IL-6 R | Reverse | TGAAGGACTCTGGCTTTGTCT |
| TNFa F | Forward | CCCTCACACTCAGATCATCTTCT |
| TNFa R | Reverse | GCTACGACGTGGGCTACAG |
| NOS2 (iNOS) F | Forward | GTTCTCAGCCCAACAATACAAGA |
| NOS2 (iNOS) R | Reverse | GTGGACGGGTCGATGTCAC |
| TBP F | Forward | AGAACAATCCAGACTAGCAGCA |
| TBP R | Reverse | GGGAACTTCACATCACAGCTC |

All sequences target mouse genes.

**Table S2 siRNA sequences used in the study.**

| Gene | Source | Identifier | Format |
|---|---|---|---|
| non-targeting control (NTC) | Horizon Discovery | D-001810-10-20 | |
| APT1 | Horizon Discovery | L-010007-00-0005 | ON-TARGET plus SMARTpool siRNA |
| PACS1 | Horizon Discovery | L-006697-01-0005 | ON-TARGET plus SMARTpool siRNA |
| YTHDF2 | Horizon Discovery | L-021009-02-0005 | ON-TARGET plus SMARTpool siRNA |
| STX12 | Horizon Discovery | L-018246-01-0005 | ON-TARGET plus SMARTpool siRNA |
| Clathrin HC | Horizon Discovery | J-004001-09-0002 | ON-TARGET plus individual siRNAs |
| Caveolin 1 | Horizon Discovery | J-003467-06-0002 | ON-TARGET plus individual siRNAs |

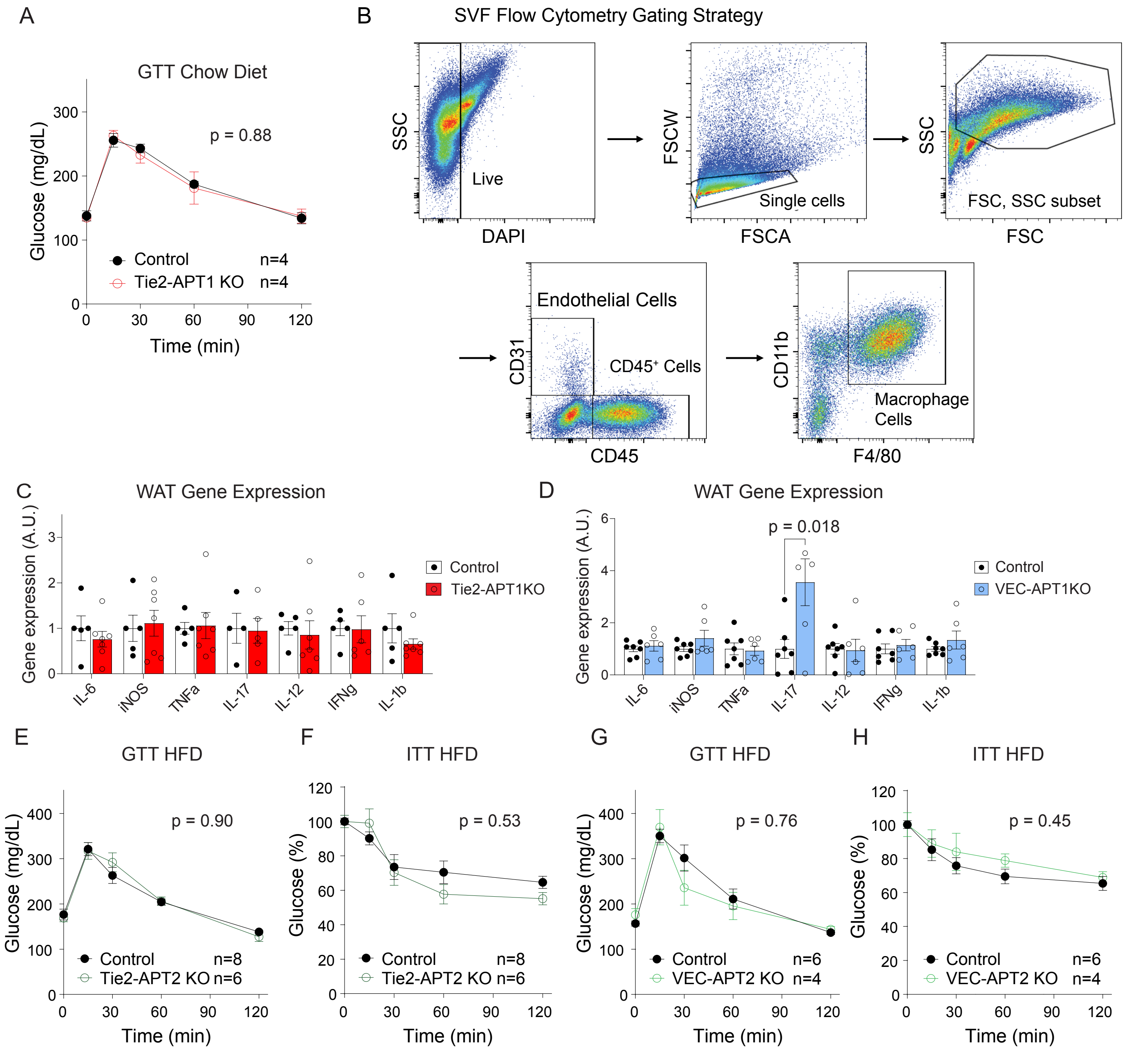


**Figure S1. Endothelial-specific APT1 knockout improves glucose tolerance in HFD-fed mice without impacts on inflammation, related to Figure 1.**

(**A**) GTTs for Tie2-APT1 KO and control male mice with chow diet feeding. (**B**) Flow cytometry gating strategy for total macrophages (CD45+CD11b+F4/80+) from the stromal vascular fraction (SVF) of white adipose tissue. (**C**) Inflammatory marker gene expression in white adipose tissue of HFD-fed Tie2-APT1 KO male mice. (**D**) Inflammatory marker gene expression in white adipose tissue of HFD-fed VEC-APT1 KO male mice. (**E-F**) GTTs (E) and ITTs (F) for Tie2 APT2 KO male mice and littermate control mice after 3 months of HFD feeding. (**G-H**) GTTs (G) and ITTs (H) for VEC-APT2 KO male mice and littermate control mice after 3 months of HFD feeding. Mice were fasted for 4–6 h prior to GTTs or ITTs. Data are expressed as mean ± SEM. Statistical analyses for genotype comparisons were performed by two-way ANOVA or by unpaired t-test with Welch's correction.

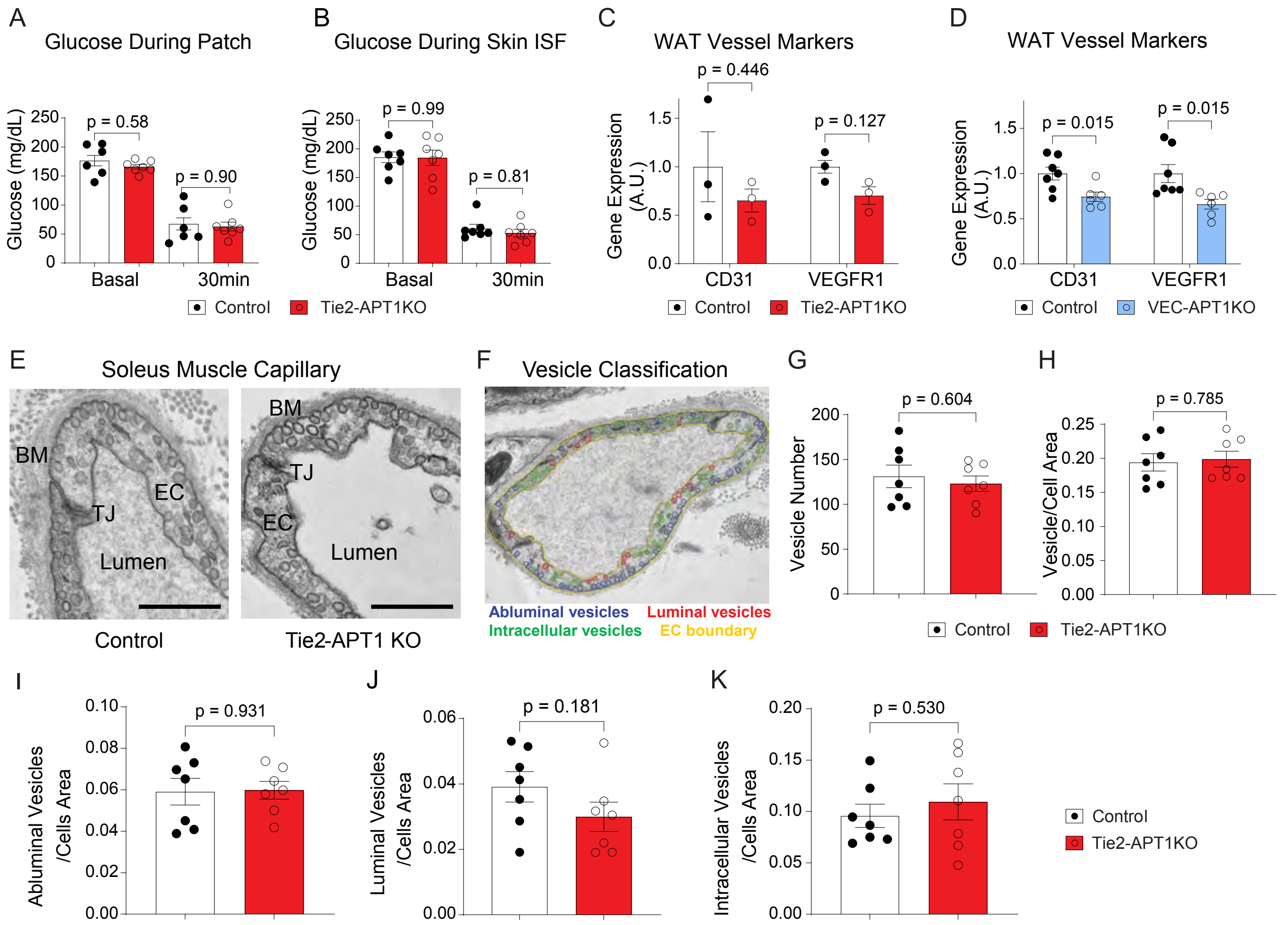


**Figure S2. APT1 deficiency does not increase vascular areas and does not affect intracellular vesicles, related to Figure 2.**

(**A-B**) Glucose levels before and 30 min after insulin injection during patch experiment (A) and skin ISF experiment (B). (**C-D**) White adipose tissue vessel marker gene expression by q-PCR. HFD-fed Tie2-APT1 KO and control mice (C). HFD-fed VEC-APT1 KO and control mice (D). (**E**) Representative TEM images of soleus muscle capillary of HFD-fed Tie2-APT1 KO and control mice. EC: endothelial cell, TJ: tight junction, BM: basement membrane. Scale = 600 nm. (**F**) Vesicle classification. Vesicles were assigned different colors based on their locations. The endothelial cell boundary was outlined in yellow. (**G-K**) TEM vesicle quantification: total vesicle number (G), vesicle density (H), and quantification of specified abluminal vesicles (I), luminal vesicles (J), and intracellular vesicles (K) in control and KO mice (2 mice/group, HFD). Data are expressed as mean ± SEM. Statistical analyses were performed by two-way ANOVA with Sidak's post hoc test or by unpaired t-test with Welch's correction.

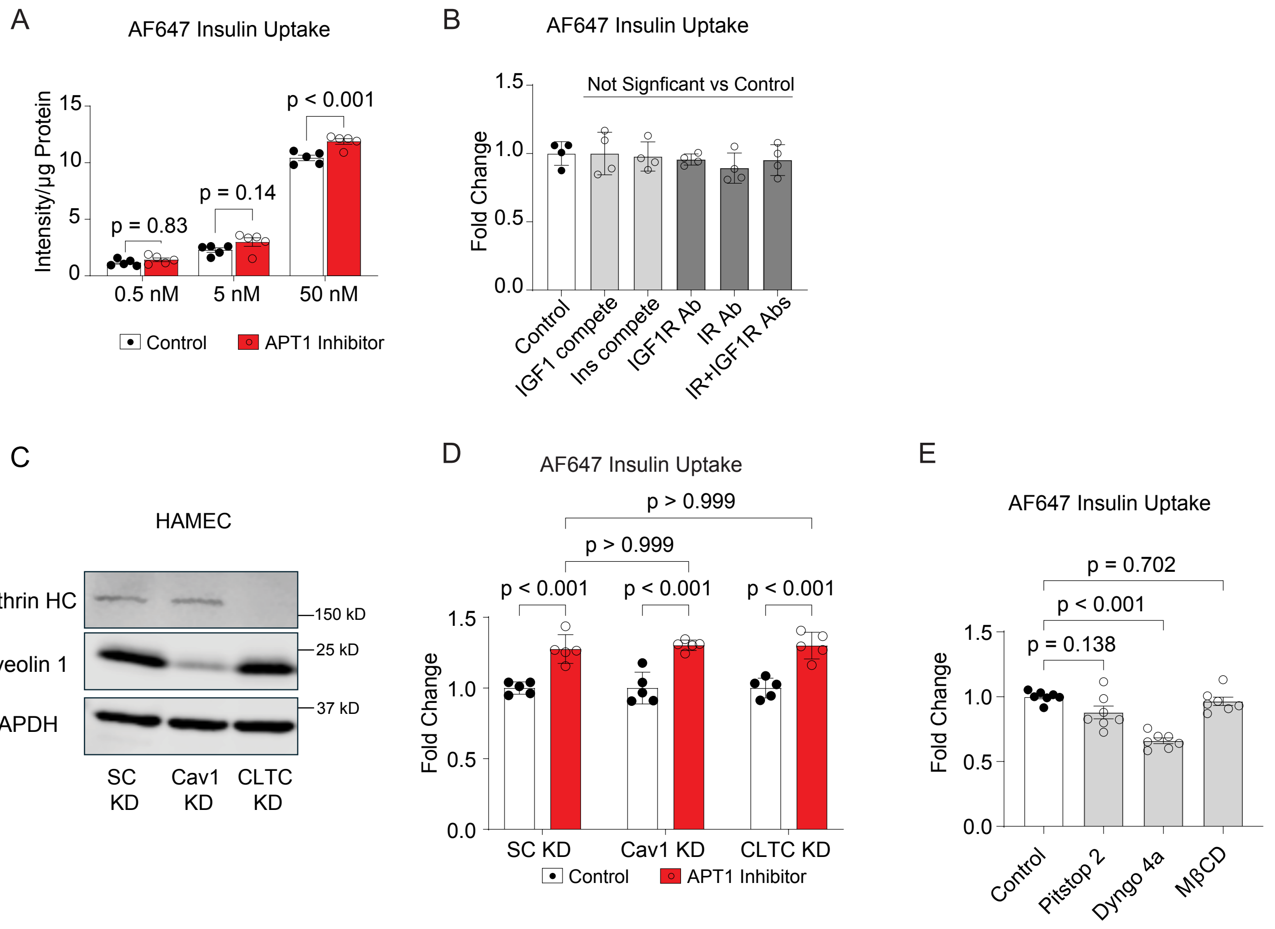


**Figure S3. APT1 selectively modulates endothelial insulin uptake through a non-canonical pathway, related to Figure 3.**

(**A**) Insulin uptake was assayed in HAMECs loaded with various concentrations of AF647 insulin in the absence or presence of the APT1 inhibitor ML348 (10 µM for 3 hrs). (**B**) AF647 insulin uptake (100 nM) in HAMECs pretreated for 5 min with control buffer, IGF1 (10 µM), native insulin (10 µM), IGF1 receptor antibody (clone 1H7, 10 µg/mL), insulin receptor antibody (clone 47-9, 10 µg/mL), or both IGF1R/IR antibodies. (**C**) Knockdown efficacy confirmed by Western blot. (**D**) AF647 insulin uptake in HAMECs with scrambled control, caveolin-1 knockdown, or clathrin knockdown, in the absence or presence of APT1 inhibitor ML348. (**E**) AF647 insulin uptake in HAMECs pretreated for 1 h with 10 µM Pitstop 2 (clathrin inhibitor), 10 µM methyl-β-cyclodextrin (MβCD), or 30 µM Dyngo-4a (dynamin inhibitor). Data are expressed as mean ± SEM. Statistical analyses were performed by two-way ANOVA with Sidak's post hoc test or one-way ANOVA with Dunnett T3 post hoc test.

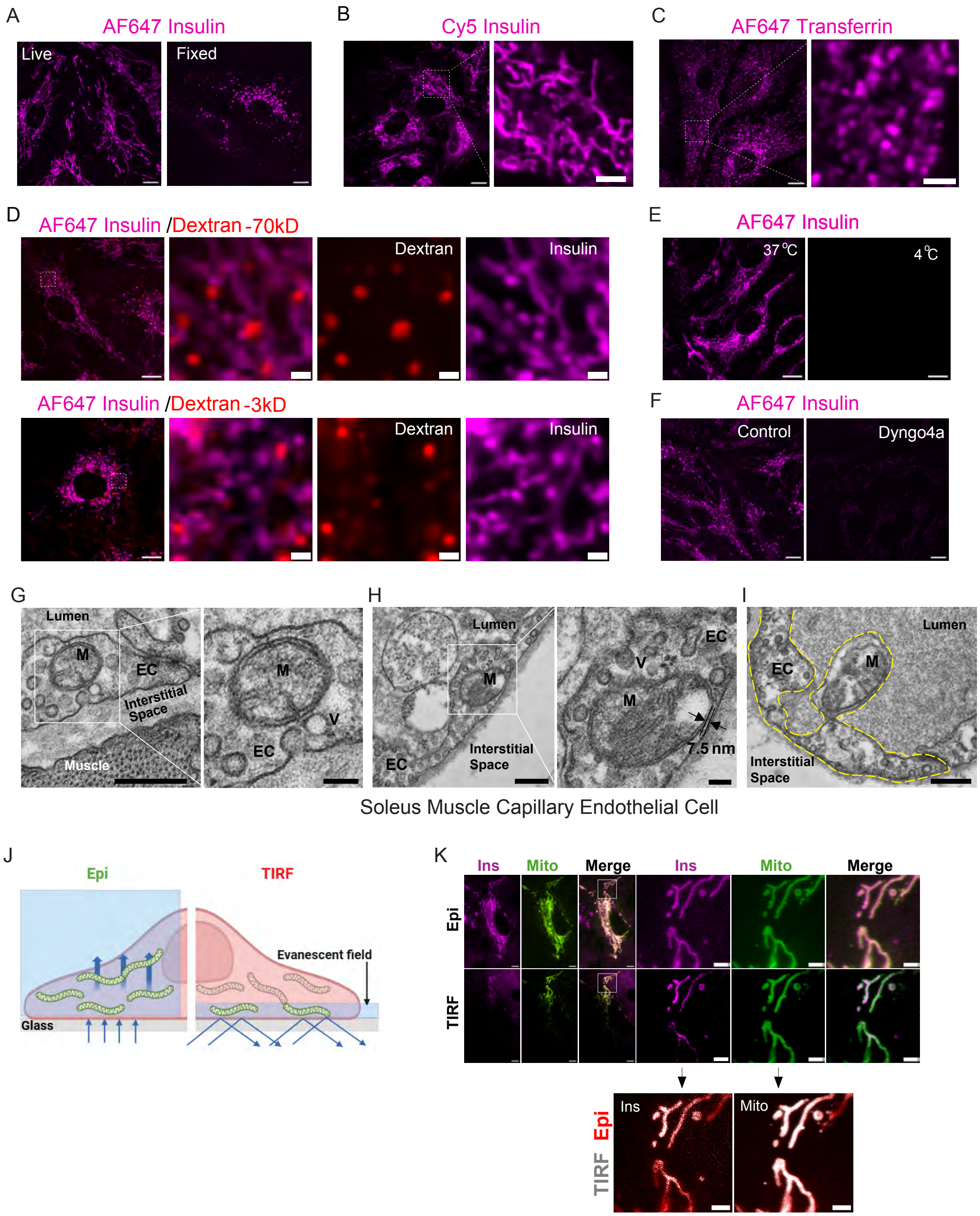


**Figure S4. Insulin uptake is involved with mitochondrial-associated tubular network in endothelial cells, related to Figure 4.**

(**A**) Representative live-cell and PFA-fixed confocal images of HAMECs treated with 50 nM AF647 insulin, scale = 10 μm. (**B**) Live-cell images of HAMECs treated with Cy5-labeled insulin (500 nM, 15min). (**C**) Live-cell images of cells treated with AF647- labeled transferrin (100 μg/mL, 15 min). (**D**) Live-cell images of HAMECs co-treated with AF647 insulin (500 nM, 15 min) and Texas Red-Dextran 70kD or 3kD (25 μg/mL, 15min). (**E**) Live-cell images of cells treated with AF647 insulin at 37°C and 4°C. (**F**) Dynamin inhibitor (Dyngo-4a, 30 μM for 10 min) effectively blocked AF647 insulin uptake. Images used same contrast setting to compare the signal differences. (**G-I**) Representative TEM images of mitochondria (M) in capillary endothelial cells (EC) of soleus muscle from HFD-fed WT mice. G: mitochondria in proximity to abluminal vesicles. H: mitochondria near the abluminal membrane (~7.5 nm distance). I: rare event of mitochondrial extrusion into the EC lumen. The cell boundary was highlighted with a yellow line. V: vesicles. (**J**) Schematic diagram illustrating differences between epifluorescence and TIRF microscopy. On the left, the incident laser (blue arrows) goes through the whole cell, thus exciting fluorescence of all mitochondria with GFP expression (green). On the right, the incident laser beam (blue arrows) is applied at a supra-critical angle that enables the total internal reflection, creating a thin evanescent zone (blue field). Mitochondria within the evanescent fields are excited and shown in green, while those outside the field are not excited and invisible. (**K**) Representative live-cell images of HAMECs treated with AF647 insulin under epifluorescence and TIRF modes. Mitochondria were labeled with MAO-GFP. A: Scale = 5 μm. B-C: Scale = 10 μm (2 μm in zoomed-in images). D: Scale = 5 μm (1 μm in zoomed-in images) E-F: Scale = 10 μm (1 μm in zoomed-in images). G-I: Scale = 400 nm (100 nm in zoomed-in images), K: Scale = 10 μm (1 μm in zoomed-in images).

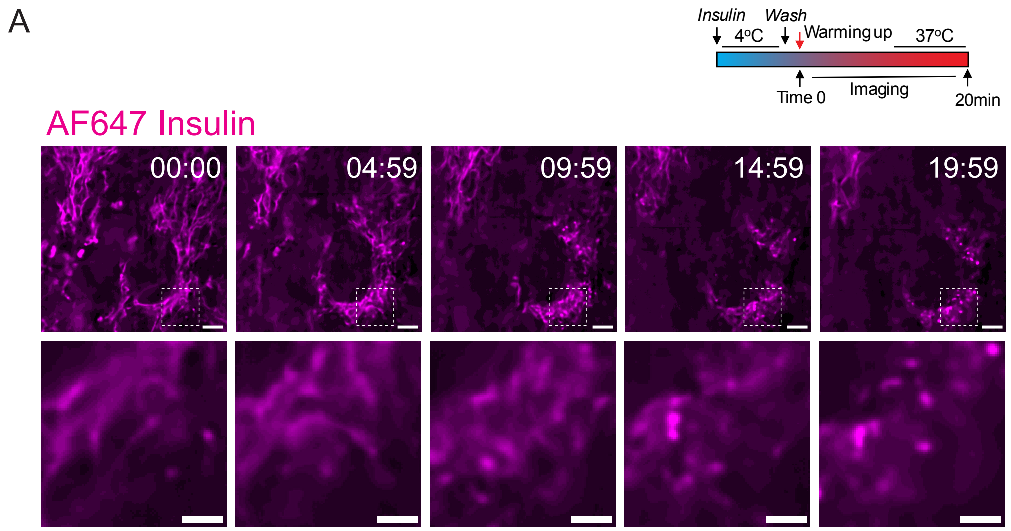


**Figure S5. Dynamics of intracellular insulin trafficking from mitochondria to late endosome, related to Figure 5.**

(**A**) Representative time-lapse images of AF647 insulin in HAMECs with timestamps indicating time after the imaging chamber was warmed up. A schematic of the insulin loading, washing, and imaging steps is shown. Scale = 5 μm (2 μm in inserts).

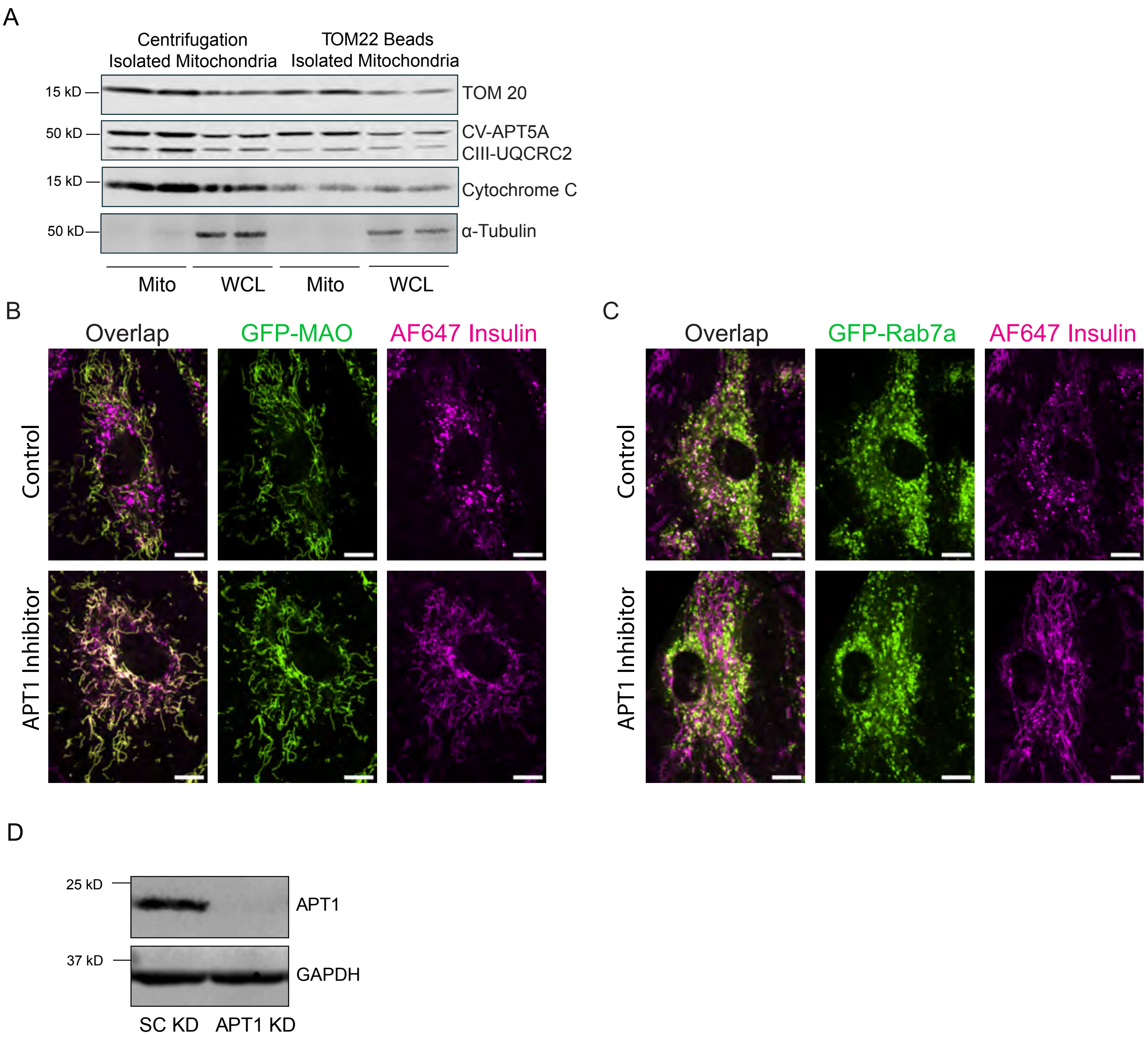


**Figure S6. APT1 inhibition does not induce morphological changes in mitochondria and Rab7a marked late endosome, related to Figure 6.**
(**A**) Western blot validation of mitochondria isolation from HAMECs using either centrifugation fractionation or TOM22 antibody bead pull-down methods. (**B**) Representative live-cell images of AF647 insulin with a mitochondrial marker (GFP-MAO) in HAMECs in the absence or presence of APT1 inhibition, scale = 10 μm. (**C**) Representative live-cell images of AF647 insulin with a late endosome marker GFP-Rab7a in HAMECs in the absence or presence of APT1 inhibition, scale = 10 μm. (**D)** APT1 knockdown was confirmed with Western blot for Figure 6C.

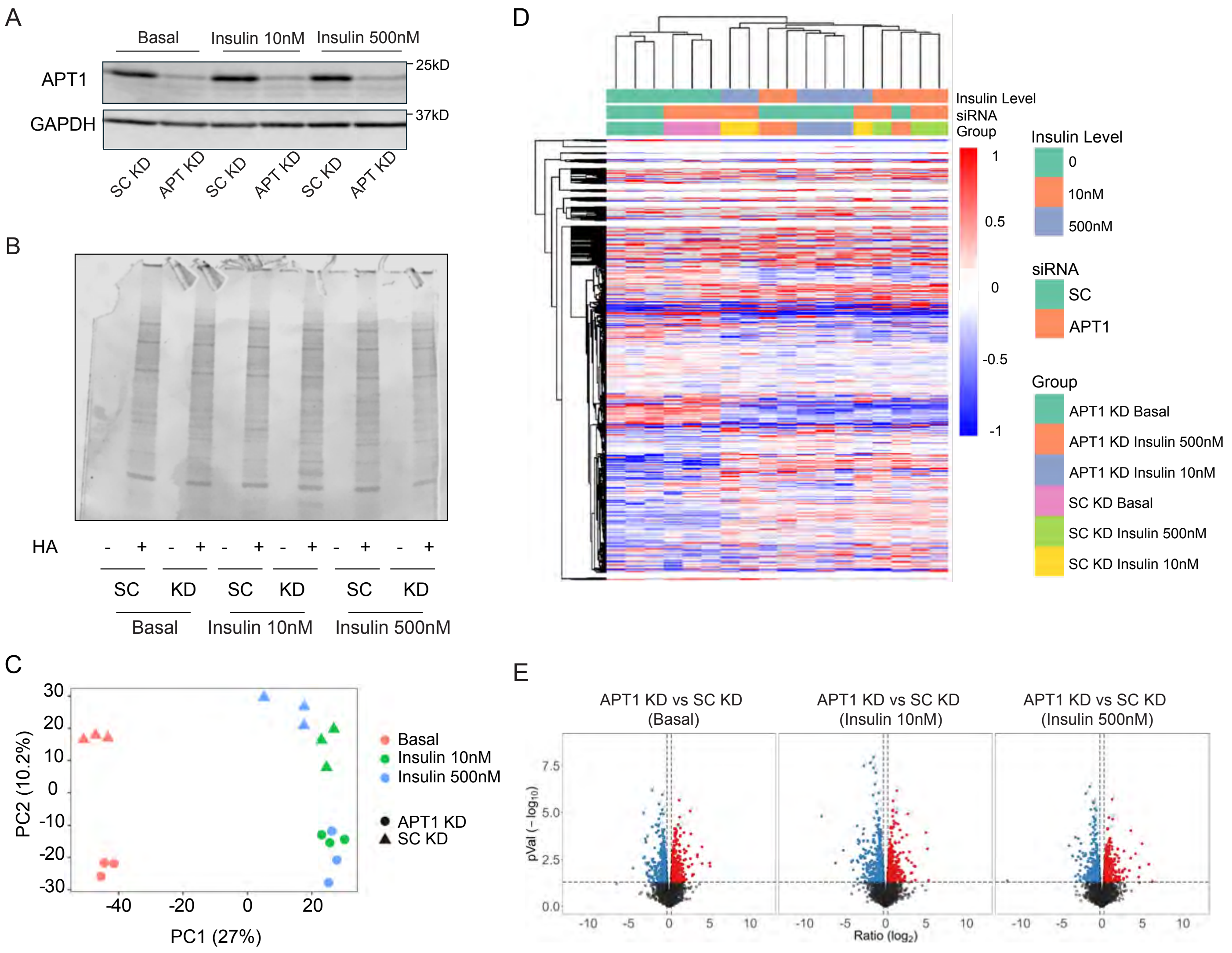


F

| Palmitoyl Proteomics Candidates \ Condition | Insulin - | | Insulin 10nM | | Insulin 500nM | |
|---|---|---|---|---|---|---|
| | KD vs SC | KD vs SC | KD vs SC | KD vs SC | KD vs SC | KD vs SC |
| | pVal | log2FC | pVal | log2FC | pVal | log2FC |
| PACS1 | 0.0021 | 0.47 | 0.0000074 | 0.87 | 0.0011 | 0.51 |
| YTHDF2 | 0.028 | 1.79 | 0.0012 | 2.95 | 0.028 | 1.79 |
| STX12 | 0.012 | 0.43 | 0.016 | 0.41 | 0.011 | 0.44 |

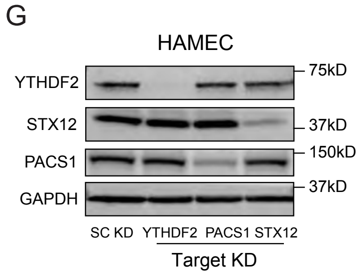


**Figure S7. Proteomic identification of APT1-dependent palmitoylation proteins, related to Figure 7.** (**A**) Validation of APT1 knockdown in HAMEC lysates used for proteomics. (**B**) Confirmation of RAC bead pulldown efficacy. HAMEC lysates were incubated with RAC beads in the presence or absence of hydroxylamine (HA). Eluted proteins were subjected to SDS-PAGE and visualized by Coomassie Blue staining. (**C**) Principal component analysis of the palmitoylation proteomics data. (**D**) Heatmap of global palmitoylation identified by proteomics. (**E**) Volcano plots of changes in palmitoylation proteomics data in each insulin treatment group. F, Summary of proteomic analysis of palmitoylation for PACS1, YTHDF2, and STX12. G, Knockdown of PACS1, YTHDF2, and Syntaxin 12 (STX12) in HAMECs confirmed by Western blot.
